# Two-photon all-optical electrophysiology for the dissection of larval zebrafish brain functional connectivity

**DOI:** 10.1101/2024.01.08.574771

**Authors:** Lapo Turrini, Pietro Ricci, Michele Sorelli, Giuseppe de Vito, Marco Marchetti, Francesco Vanzi, Francesco S. Pavone

## Abstract

One of the most audacious goals of modern neuroscience is unraveling the complex web of causal relations underlying the activity of neuronal populations on a whole-brain scale. This endeavor, prohibitive just a couple of decades ago, has recently become within reach owing to the advancements in optical methods and the advent of genetically encoded indicators/actuators. These techniques, applied to the translucent larval zebrafish have enabled recording and manipulation of the activity of extensive neuronal populations spanning the entire vertebrate brain. Here, we present the conception of a custom two-photon optical system, coupling light-sheet imaging and 3D excitation with acousto-optic deflectors for simultaneous high-speed volumetric recording and optogenetic stimulation. By employing a zebrafish line with pan-neuronal expression of both the calcium reporter GCaMP6s and the red-shifted opsin ReaChR, we implemented a crosstalk-free, noninvasive all-optical approach and applied it to the reconstruction of the functional connectivity of the left habenula.

## Introduction

Understanding the functional connectivity of intricate networks within the brain is a fundamental goal towards unraveling the complexities of neural processes. This longtime focal point in neuroscience requires methodologies to trigger and capture neuronal activity in an intact organism. Critical insights into the complex interplay among large populations of neurons have been provided by electroencephalography and functional magnetic resonance imaging^1-4^. Those gold standard methods, however, do provide a noninvasive means to detect neuronal activity, but with limited spatial/temporal resolution (not allowing measurement of single neuron activity) and lack equally noninvasive possibilities to precisely elicit and control it. Therefore, it is evident that deciphering how individual neurons communicate to shape functional neural circuits on a whole-organ scale demands further technological advances. Over the last few decades, with the advent of optogenetics^5^ and the widespread adoption of genetically encoded fluorescent indicators^6^, all-optical methods have gained traction for their ability to simultaneously monitor and manipulate the activity of multiple neurons within the intact brain^7-10^.

In this framework, the ever-increasing use of the tiny and translucent zebrafish larva as an animal model^11^, has provided momentum for the development and enhancement of optical technologies aimed at imaging and controlling neuronal activity with light at high spatio-temporal resolution^12-14^. On the imaging side, previous high-resolution all-optical investigations on zebrafish have made use of two-photon (2P) point scanning methods^15-17^ or, more rarely, of one-photon (1P) excitation light-sheet fluorescence microscopy (LSFM)^18^. Compared to point scanning approaches, LSFM^19^, allowing parallelization of the detection process within each frame, enables concurrent high spatio-temporal resolution and extensive volumetric imaging. However, the use of visible excitation in 1P LSFM represents an undesired source of strong visual stimulation for the larva, often requiring tailored excitation strategies at least to prevent direct illumination of the eyes^20^.

On the photostimulation side, most advanced all-optical setups typically adopt parallel illumination approaches, making use of spatial light modulators (SLM) or digital micromirror devices (DMD) to generate multiple simultaneous holographic spots of excitation^16,21-23^. In computer-generated holography, the input laser power is subdivided among the various spots, resulting in increasing energies released on the specimen as the number of effective targets rises and, consequently, in increasing probability of photodamage^24^. Conversely, scan-based sequential stimulation allows a fraction of the power needed by parallel approaches to be deposited at any time on the sample, regardless of the number of targets. As a drawback, however, scan-based methods typically employ mechanical moving parts which constrain the stimulation sequence speed and thus the maximum temporal resolution achievable. An exception is represented by acousto-optic deflectors (AODs)^25^ which are not affected by mechanical inertia and thus enable discontinuous three-dimensional trajectories with constant repositioning time. In particular, featuring an ultrashort access time (μs range), AODs represent the scanning technology that gets closest to parallel illumination performance. Indeed, AODs enable quasi-simultaneous three-dimensional targeting of multiple spots, yet keeping low the global energy delivered. However, despite their extensive use for fast 3D imaging^26-30^, these devices have been rarely employed for photostimulation so far^31-35^.

In this work, we present an all-optical setup consisting of a light-sheet microscope and a light-targeting system equipped with AODs, both employing nonlinear excitation, which enable high spatio-temporal resolution volumetric imaging of the larval zebrafish brain along with concomitant three-dimensional optogenetic stimulation. Using a double transgenic line pan-neuronally expressing both the green fluorescent calcium indicator GCaMP6s^36^ and the red-shifted light-gated cation channel ReaChR^37^, we demonstrate a crosstalk-free experimental approach for all-optical investigation of brain circuitries. Leveraging two-photon excitation and the inertia-free light targeting capabilities of AODs, we validated the system functionality by reconstructing the efferent functional connectivity of the left habenula, a cerebral nucleus mainly composed of excitatory neurons, linking forebrain and midbrain structures.

## Results

### A crosstalk-free approach for two-photon all-optical investigations in zebrafish larvae

In order to explore brain functional connectivity in zebrafish larvae, we have devised an integrated all-optical two-photon system capable of simultaneously recording and stimulating neuronal activity. The setup, as depicted in Figure 1a, consists of a light-sheet fluorescence microscope and a light-targeting system, specifically designed for fast whole-brain calcium imaging and 3D optogenetic stimulation, respectively. Both optical paths employ pulsed near-infrared (NIR) laser sources for 2P excitation^38^. The 2P LSFM module employing digitally scanned mode, double-sided illumination, control of excitation light polarization^39^ and remote focusing of the detection objective, is capable of recording the entire larval brain (400×800×200 μm^3^) at volumetric rates up to 5 Hz^40^. On the other hand, the light-targeting system incorporates two couples of acousto-optic deflectors to move the excitation focus to arbitrary locations inside a 100×100×100 μm^3^ volume, guaranteeing constant access time (10 μs) independently from the relative distance between sequentially illuminated points, and equal energy delivered independently from the number of targets^35^.

**Figure 1.**
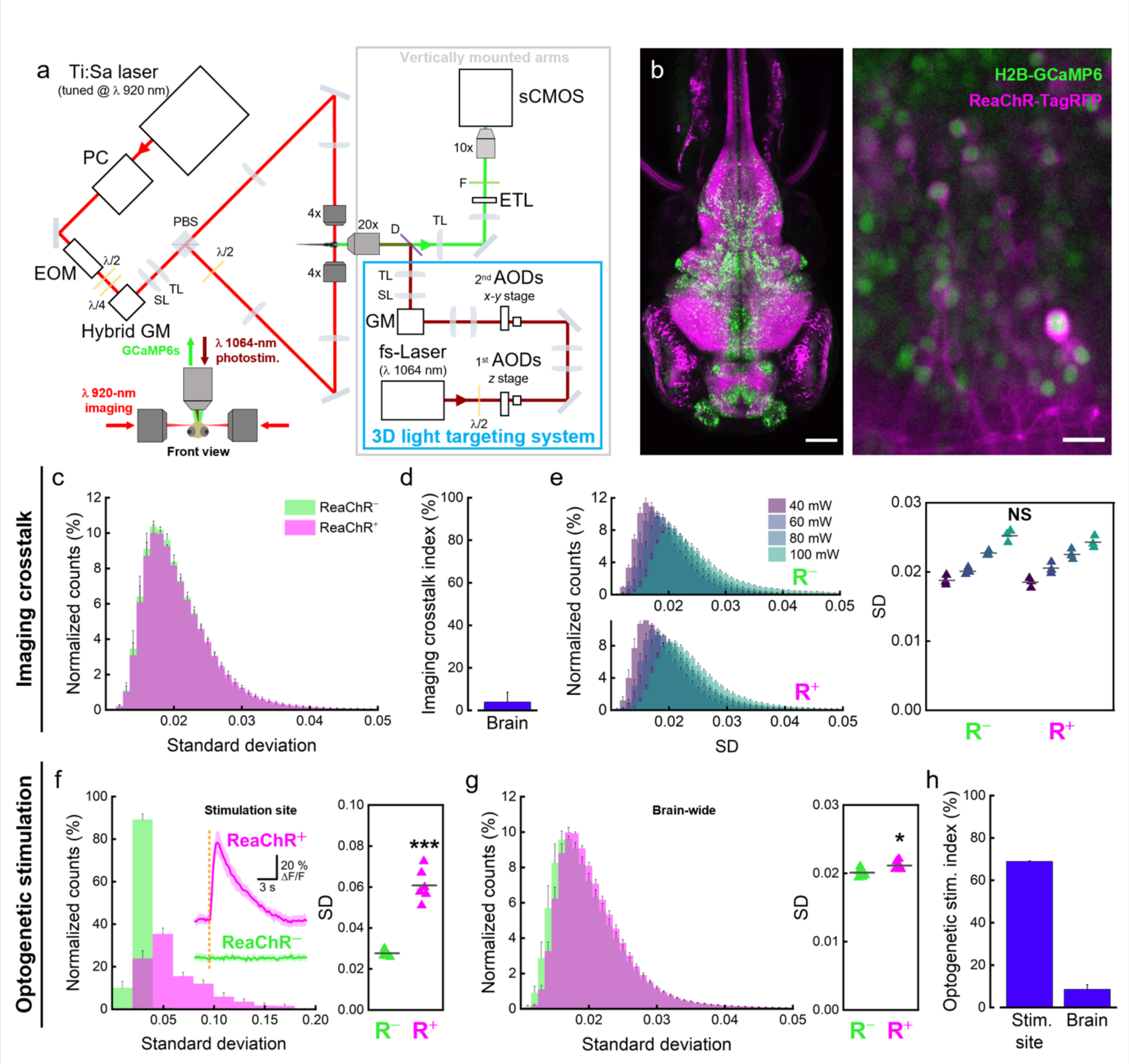
A crosstalk-free experimental paradigm for two-photon all-optical electrophysiology of the larval zebrafish brain. **a.** Schematic representation of the optical setup devised for simultaneous whole-brain imaging and optogenetic stimulation. Red line indicates the 920 nm excitation path employed for light-sheet imaging, green indicates GCaMP6s fluorescence in the imaging detection path, darker red indicates the 1064 nm excitation of the 3D light-targeting system used for stimulation. Arrows at the beginning of each optical path indicate the direction of light propagation. PC, pulse compressor; EOM, electro-optical modulator; λ/2, half-wave plate; λ/4, quarter-wave plate; GM, galvo mirrors; SL, scan lens; TL, tube lens; PBS, polarizing beam splitter; D, dichroic mirror; ETL, electrically tunable lens; F, fluorescence filter; AODs, acousto-optic deflectors. See Methods for details. Inset at the bottom represents the front view of a zebrafish larva inside the imaging chamber. The two opposite illumination objectives are employed to focus the 920-nm virtual light-sheet (red) inside the larval brain. The detection objective is used both for GCaMP6s fluorescence (green) collection and for focusing the 1064-nm excitation laser (darker red) for optogenetic stimulation. **b.** Double transgenic zebrafish line for all-optical electrophysiology. **Left**, maximum intensity projection of a confocal stack encompassing the entire head of a 5 dpf *Tg(elavl3:H2B-GCaMP6s; elavl3:ReaCHR-TagRFP)* larva (see also Supplementary Movie 1). **Right**, higher magnification confocal image of reticulospinal neurons showing the nuclear expression of the calcium reporter GCaMP6s (green) and membrane localization of the fusion protein ReaChR-TagRFP (magenta). Scale bars: 100 μm (left), 10 μm (right). **c-d-e.** Quantification of crosstalk activation of ReaChR channels during 5 minutes of light-sheet imaging. N = 3 ReaChR^+^ and 3 ReaChR^−^ larvae. **c.** Average normalized distributions of the SD values of voxels composing the larval brain over 5 minutes of continuous light-sheet imaging at λex 920 nm and power 60 mW for ReaChR^+^ and ReaChR^−^ larvae. **d.** Imaging crosstalk index, calculated as the percentage Hellinger distance (see Methods for equation) between the average normalized distributions shown in Figure 1c. **e. Left**, Average normalized distributions of the SD values of voxels composing the larval brain over 5 minutes of continuous light-sheet imaging at increasing laser power (40, 60, 80 and 100 mW) for ReaChR^−^ (R^−^, upper) and ReaChR^+^ (R^+^, lower) larvae. **Right**, median SD values calculated from individual SD distributions of brain-wide voxels of R^−^ and R^+^ larvae. Statistical comparisons (two-way ANOVA and post-hoc Tukey’s test) highlight no statistical significance (*P*>0.05) between SD values of R^−^ and R^+^ larvae exposed to the same imaging power and between SD values of R^+^ or R^−^ larvae exposed to 40 and 60 mW power. **f-g-h.** Quantification of the effect of the optogenetic stimulation on neuronal activity. **f. Inset,** nuclear GCaMP6s signals induced by 500 ms stimulation at 1064 nm while performing whole-brain light-sheet imaging at 920 nm. Larvae expressing ReaChR and GCaMP6s show stimulation-induced calcium transients (magenta trace) whereas larvae expressing only GCaMP6s (green trace) do not. Orange dashed line indicates the stimulation event. N = 3 ReaChR^+^, 3 ReaChR^−^ larvae; n = 5 calcium transients per larva. Data are presented as mean ± sd. **Left,** average normalized distributions of the SD values of voxels inside the stimulation site for ReaChR^+^ and ReaChR^−^ larvae over 100 s of whole-brain imaging during which 5 optogenetic stimulation were delivered. N = 6 ReaChR^+^, 6 ReaChR^−^ larvae. **Right,** median SD values calculated from individual SD distributions of stimulation site voxels for R^−^ and R^+^ larvae. *** *P* < 0.0001, unpaired *t* test. **g. Left,** average normalized distributions of the SD values of brain-wide voxels for ReaChR^+^ and ReaChR^−^ larvae in the same condition as in panel f. Right, median SD values calculated from individual SD distributions of brain-wide voxels for R^−^ and R^+^ larvae. * *P* = 0.0218, unpaired *t* test. **h.** Optogenetic stimulation index, percentage Hellinger distance between the average normalized distributions shown in panel f (stim. site) and g (brain), respectively. Error bars of average distributions indicate the sem. Error bars of imaging crosstalk and optogenetic stimulation indices were calculated according to uncertainty propagation theory (see Methods for details). Gray horizontal bars in panels e, f and g represent intragroup mean values.

In order to perform simultaneous recording and stimulation of neuronal activity, we employed the pan-neuronal *Tg(elavl3:H2B-GCaMP6; elavl3:ReaChR-TagRFP)* zebrafish line (Figure 1b, left, Supplementary Movie 1). Larvae of this double transgenic line express the green fluorescent calcium indicator GCaMP6s inside neuronal nuclei and the red-shifted light-gated cation channel ReaChR (as a fusion protein with the red fluorescent protein TagRFP) on neuronal membranes (Figure 1b, right).

We first investigated the possible presence of crosstalk activation of ReaChR (which for simplicity will be hereafter written as R) channels due to the excitation wavelength used for functional imaging. To this end, light-sheet imaging of both double transgenic larvae (R^+^) and GCaMP6s expressing larvae (R^−^, lacking the light-gated channel) was performed for 5 minutes (volumetric rate: 2.5 Hz, λex: 920 nm, laser power at the sample: 60 mW). To evaluate the level of neuronal activity, we computed the standard deviation (SD) over time of each voxel belonging to the brain. Between the two groups, no major differences could be observed in the average SD distributions computed over a 5-minute exposure to the imaging laser (Figure 1c). Indeed, the resulting imaging crosstalk index (calculated as the Hellinger distance between the two average distributions, see Methods for details) was extremely low (3.9% ± 4.5%; Figure 1d). However, since crosstalk activation of light-gated channels by a spurious wavelength is typically power-dependent^41^, we then investigated whether higher powers of the laser used for imaging could induce a significant effect on R^+^ larvae. Figure 1e shows the average SD distributions obtained from R^+^ and R^−^ larvae at imaging power ranging from 40 to 100 mW. Despite higher laser powers producing a shift of the distributions towards higher SD values, this equally affects the neuronal activity of both R^+^ and R^−^ larvae (Supplementary Figure 1a). The differences between the median values of the SD distributions of the two groups (R^+^ and R^−^) at the same imaging power are not statistically significant (Figure 1e, right) and, indeed, the imaging crosstalk index remains essentially constant in the power range tested (Supplementary Figure 1b). After having demonstrated the absence of crosstalk activation of ReaChR channels upon 2P light-sheet scanning, we investigated the capability of our AOD-based photostimulation system to effectively induce optogenetic activation of targeted neurons. For this purpose, we stimulated R^+^ larvae at 1064 nm (laser power at the sample: 30 mW, stimulation volume: 30×30×30 μm^3^) while simultaneously recording whole-brain neuronal activity with light-sheet imaging.

Larvae expressing the opsin showed strong and consistent calcium transients evoked in the stimulation site (Figure 1f, inset). Conversely, stimulating R^−^ larvae did not result in any detectable response (Figure 1f, inset). We quantified the effect of the optogenetic stimulation by computing the distributions of SD values of the voxels inside the stimulation site for R^+^ and R^−^ larvae (Figure 1f, left). 1064 nm stimulation induced statistically significant optogenetic activation of opsin-expressing neurons in R^+^ larvae (R^−^ = 0.0277 ± 0.0017, R^+^ = 0.0608 ± 0.0077; Figure 1f, right). Despite the small stimulation volume with respect to the entire brain size, the effect of the photostimulation was noticeable also in the whole-brain SD distribution, where R^+^ larvae showed a peak slightly shifted towards higher SD values (Figure 1g, left) which produced a significantly higher average SD (R^−^ = 0.0201 ± 0.0007, R^+^ = 0.0211 ± 0.0006; Figure 1g, right). This noticeable difference is due to the high-amplitude calcium transients evoked by the stimulation and by the activation of the neuronal population synaptically downstream to the stimulation site. Figure 1h shows the optogenetic activation index (calculated as the Hellinger distance between the two average distributions, see Methods for details) for the stimulation site and the entire brain (stimulation site = 68.9% ± 0.3%, brain = 8.6% ± 2.1%). In order to rule out any possible spurious activation effect not related to the optogenetic excitation of ReaChR channels (e.g., sensorial perception of the laser stimulus), we also compared the SD distributions of ReaChR^−^ larvae only subjected to imaging (R^−^i) and to simultaneous imaging and stimulation (R^−^i+s). The analysis highlighted no statistically significant effects of the photostimulation in absence of opsin expression both in the stimulus site (Supplementary Figure 2a) and at a brain-wide level (Supplementary Figure 2b).

### Characterization of calcium transients evoked by 3D optogenetic stimulation

After assessing the absence of opsin crosstalk activation upon light-sheet imaging and verifying the capability of our system to consistently induce optogenetic activation of ReaChR^+^ neurons, we performed a characterization of the neuronal responses aiming at identifying optimal stimulation parameters. We decided to target the stimulation at an easily recognizable cerebral nucleus mainly composed of excitatory neurons. Neurons having their soma inside the habenulae express *vesicular glutamate transporter 2* (VGLUT2, also known as *SLC17A6*), representing a coherent group of excitatory glutamatergic neurons^42,43^. We therefore directed the stimulation onto the left habenula, an anatomically segregated nucleus, part of the dorsal-diencephalic conduction system^44^. We adopted a stimulation volume of 50×50×50 μm^3^, sufficient to cover the entire habenula (Figure 2a).

**Figure 2.**
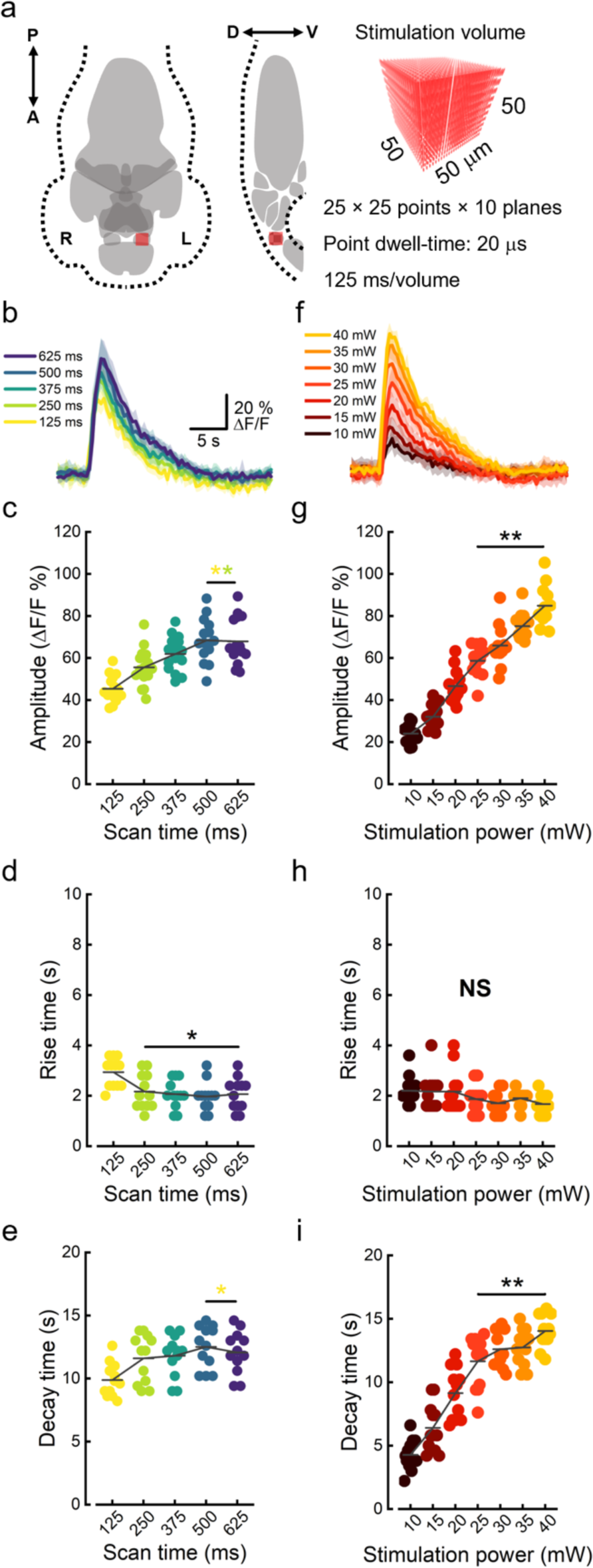
Characterization of the neuronal activation in response to optogenetic stimulation. **a.** Schematic representation of the larval brain in dorso-ventral projection (left) and midline sagittal section (right). The brain is segmented into ten gray anatomical districts used for further analysis in Figures 3 and 4. In the sagittal section, habenulae are depicted with a checkered pattern since they would not be included in the represented midline plane. Red squares indicate the 50×50×50 μm^3^ stimulation volume to scale with the brain and located over the stimulation site (left habenula). The illumination spot (represented with the excitation PSF dimensions in the rendering on right) was discontinuously scanned to 6250 different locations inside the stimulation volume. With a point every 2 μm, along 10 planes displaced every 5 μm, and a dwell-time of 20 μs, a complete iteration took 125 ms. A, anterior; P, posterior; D, dorsal; V, ventral; R, right hemisphere; L, left hemisphere. **b.** Optogenetically evoked calcium transients as a function of scan time (stimulation duration). **c-d-e.** Amplitude (**c**), rise (**d**) and decay time (**e**) of stimulation-evoked calcium transients as a function of scan time. Color code as in panel b. (**c**) ** *P* < 0.00304, (**d**) * *P* < 0.02575, (**e**) * *P* < 0.01798, one way ANOVA and post-hoc Tukey’s test. Colored asterisks indicate with respect to which scan time data are significantly different. **f.** Optogenetically evoked calcium transients as a function of stimulation power. Same scale as in panel b. **g-h-i.** Amplitude (**g**), rise (**h**) and decay time (**i**) of evoked calcium transients as a function of stimulation power. Data are color-coded as in panel f. (**g**) ** *P* < 0.00431, (**h**) NS *P* > 0.2377, (**i**) ** *P* < 0.00589, one way ANOVA and post-hoc Tukey’s test. Data in panels b and f are presented as mean ± sd. Data in panels c, d, e, g, h, i represent quantities measured from individual calcium transients. Gray horizontal bars in panels c, d, e, g, h, i represent intragroup mean values. N = 4 ReaChR^+^ larvae, n = 3 calcium transients per larva.

This volume was populated with 6250 points distributed over 10 z-planes (z step: 5 μm). With a point dwell-time of 20 μs, a complete cycle over all the points in the volume took only 125 ms. We first characterized the calcium transients as a function of the stimulation duration (scan time, Figure 2b), in a range from 125 to 625 ms (1 to 5 iterations over the volume). Figure 2c shows the amplitude of calcium peaks as a function of the scan time. Increasing scan durations produced a progressive increase in peak amplitude until reaching a plateau between 4 and 5 volume cycles (scan time 500- 625 ms). From a kinetic point of view, increasing scan durations led to a significant decrease of the rise time of calcium transients (Figure 2d). Also, the decay time of calcium transients showed a progressive increase at higher scan times (Figure 2e). We also characterized the neuronal response as a function of the 1064 nm excitation power (ranging from 10 to 40 mW, Figure 2f). The amplitude of calcium transients showed a strong linear dependence on the stimulation power (Figure 2g, R^2^: 0.89). While the rise time did not seem to be affected by the laser intensity (Figure 2h), the decay time showed a strong linear proportionality (Figure 2i, R^2^: 0.82). The duration of calcium transients (Supplementary Figure 3a,b) instead increased along with the stimulation power while not being significantly affected by the scan time. Given the small variation in rise time, in both cases the overall duration of the calcium transient was largely determined by the decay time trend.

### Whole-brain functional circuitry of left habenular nucleus

After this initial technical validation, we employed our all-optical setup to identify cerebral regions functionally linked to the left habenular nucleus. To this end, we designed the following stimulation protocol (Figure 3a). For each zebrafish larva, we performed six trials consisting of 5 optogenetic stimuli (interstimulus interval: 16 s) during simultaneous whole-brain light-sheet imaging. Based on the characterization performed, we adopted a stimulus duration of 500 ms (4 complete consecutive iterations over the 50×50×50 μm^3^ volume, point density: 0.5 point/μm) and a laser power of 30 mW, to maximize the neuronal response while keeping low the laser intensity (Supplementary Movie 2). First, we evaluated the brain voxel activation probability in response to the optogenetic stimulation of the left habenula. Figure 3b shows different projections of the whole-brain average activation probability map (Supplementary Movies 3 and 4). Left habenula (LHb), which was the site of the stimulation, predictably showed the highest activation probability values. Besides the LHb, an unpaired nucleus located at the ventral midline of the posterior midbrain showed increased activation probability with respect to the surrounding tissue. Then, we segmented the entire larval brain into ten different anatomical regions according to structural boundaries (Figure 3c, left). By extracting the average activation probability from each region (Figure 3c, right), we found the deep midbrain district to correspond to the interpeduncular nucleus (IPN). IPN is a renown integrative center and relay station within the limbic system which receives habenular afferences traveling through the fasciculus retroflexus^45-47^. Figure 3d shows the average normalized activation probability distributions for voxels inside LHb, IPN and right habenula (RHb, as the region with the higher activation probability after LHb and IPN). Neurons of LHb and IPN show activation probabilities as high as 100% and 51%, respectively. Notably, despite LHb presenting higher activation probabilities compared to IPN across larvae, higher LHb probabilities do not necessarily correspond to higher IPN probabilities (Figure 3e). Figure 3f shows the mean ΔF/F signals obtained from the LHb (blue) and IPN (yellow) regions during a stimulation trial. LHb consistently responds to the photostimulation with high-amplitude calcium transients. IPN shows lower amplitude activations (about 1/10 of LHb), yet reproducibly following the pace induced by the LHb stimulation (as also visible in Supplementary Movie 2). The coherence between these time traces is confirmed by their cross power spectrum, showing highest density around the optogenetic trigger rate (1/16 Hz, Figure 3g). As a comparison, Supplementary Figure 4 shows the cross power spectral density of the LHb and RHb activities, where null to low coupling levels emerge.

**Figure 3.**
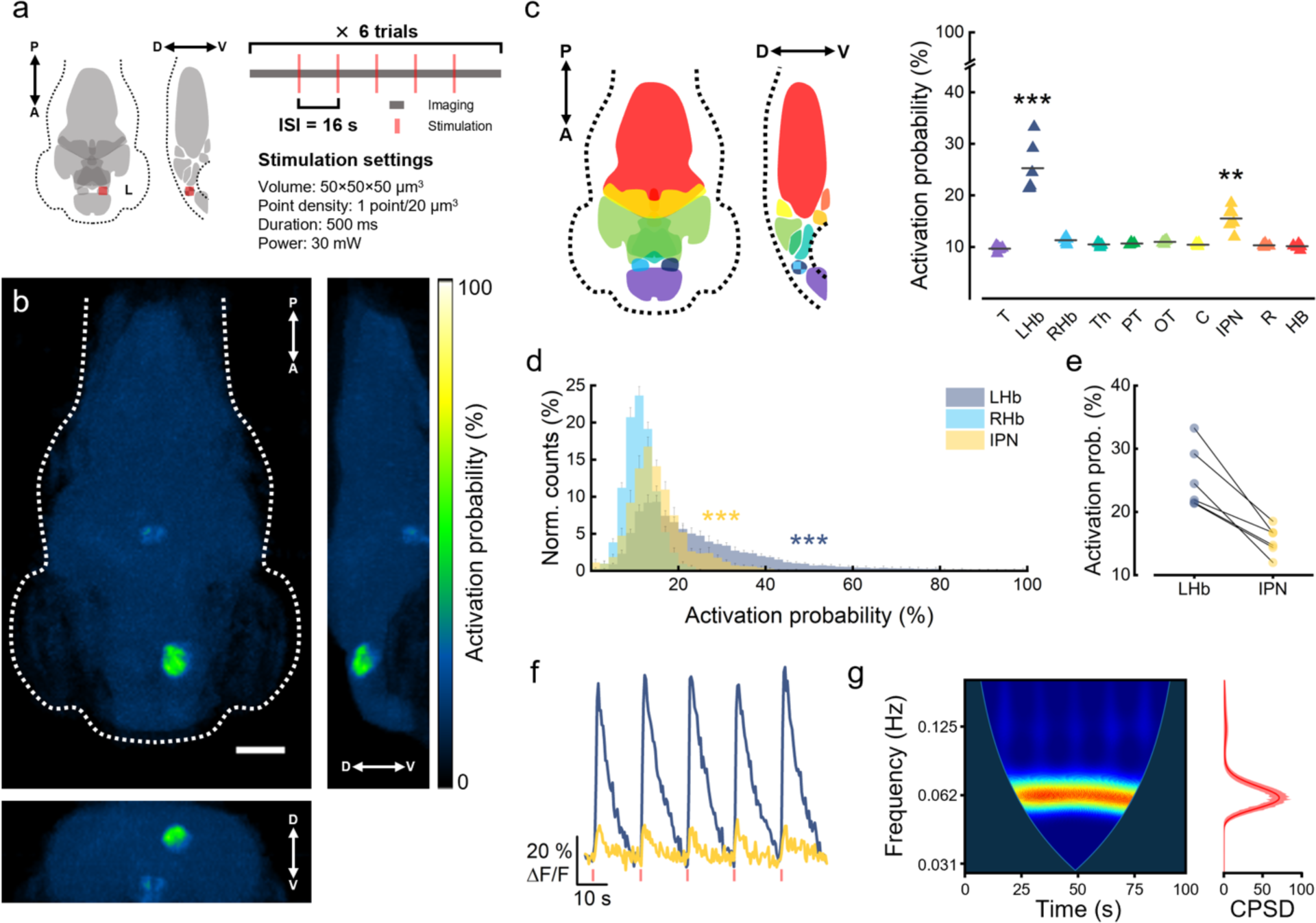
Brain-wide activation probability in response to optogenetic stimulation of the left habenula. **a.** Scheme of the stimulation protocol adopted for the investigation of habenular functional connectivity. The protocol consists of 6 trials, each composed of 100 s of whole-brain imaging (dark gray) during which 5 stimuli (red) are delivered with an interstimulus interval (ISI) of 16 s (see also Supplementary Movie 2). The stimulation volume (50×50×50 μm^3^) is centered in *xyz* on the left habenula and discontinuously scanned with 6250 focal points (scan time: 500 ms; 1064 nm power: 30 mW). A, anterior; P, posterior; D, dorsal; V, ventral; L, left hemisphere. **b.** Activation probability maps showing the probability of each brain voxel to exceed a 3-SD threshold during stimulation with respect to pre-stimulus levels. Maps are maximum intensity projections of the average 3D map (Supplementary Movie 3) along the transverse (left), sagittal (right) and coronal (bottom) axis. N = 6 ReaChR^+^ larvae, n = 30 stimulations per larva. Scale bar, 100 μm. **c.** Activation probability of different regions composing the larval brain. Median probability values are obtained from individual probability maps (Supplementary Movie 4). Regions are color-coded as in the schematic representation on the left. In the sagittal section, habenulae are depicted with a checkered pattern since they would not be included in the represented midline plane. T, telencephalon; LHb, left habenula; RHb, right habenula; Th, thalamus; PT, pretectum; OT, optic tectum; C, cerebellum; IPN, interpeduncular nucleus; R, raphe; HB, hindbrain. Gray horizontal bars represent intragroup mean values. *** *P* < 0.0001, ** *P* < 0.0059, one way ANOVA and post-hoc Tukey’s test. **d.** Average normalized distributions of activation probability values extracted from voxels inside LHb, IPN and RHb. *** *P* < 0.0001, KS test with Bonferroni correction (α = 0.01667). N = 6 ReaChR^+^larvae. **e.** Median activation probability extracted from LHb and IPN of individual larvae. Lines connect symbols belonging to the same individual. Color code as in panel c. N = 6 ReaChR^+^ larvae. **f.** Mean ΔF/F activity traces extracted from the LHb (blue) and IPN (yellow) during an individual trial of the stimulation protocol. At a glance we can observe the perfect match between the response of the two regions. N = 1 ReaChR^+^larva. See also Supplementary Movie 2. **g. Left,** wavelet cross power spectral density (CPSD) of the two time-traces reported in panel f. Warmer colors indicate higher spectral density. **Right,** time-averaged wavelet CPSD. The CPSD peak is centered at 0.0625 Hz, which corresponds to the stimulation period (16 s).

We then examined whole-brain functional connectivity during optogenetic stimulation. To this aim, we first extracted the neuronal activity from previously segmented brain regions. Figure 4a shows, as an example, the heatmap of the neuronal activity over time during a single stimulation trial. LHb and IPN are apparently the only two regions following the photostimulation trigger (red vertical bars at the bottom). This result is confirmed and generalized by the chord diagram presented in Figure 4b. This chart presents the average all-against-all correlation between the neuronal activity of different brain regions. LHb and IPN are the two anatomical districts that show the strongest functional connectivity during stimulation (Pearson’s correlation coefficient = 0.605 ± 0.079).

**Figure 4.**
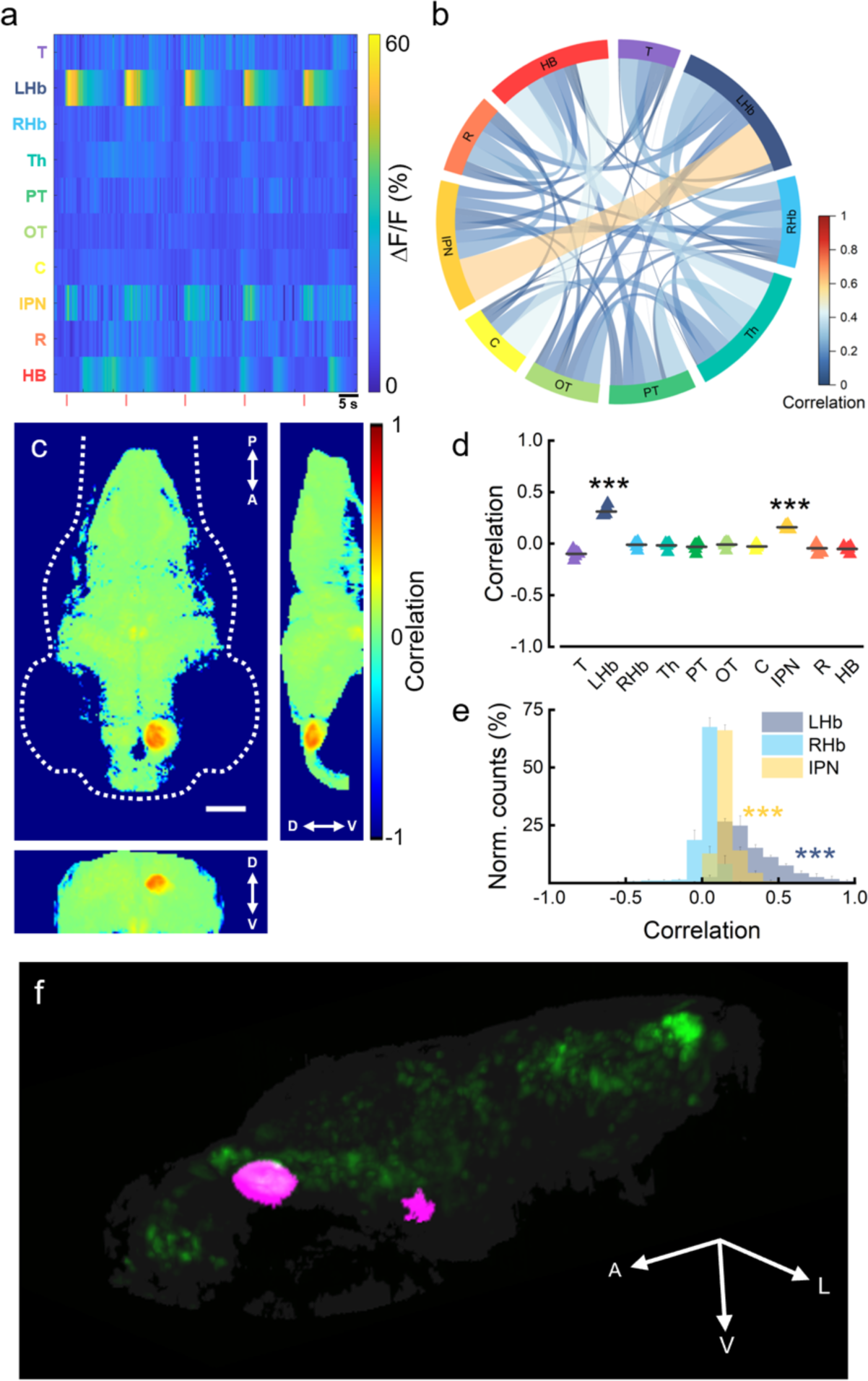
Investigation of the efferent functional connectivity of the left habenular nucleus. **a.** Heatmap showing in color-code the neuronal activity (*ΔF/F*) over time of each of the ten brain regions (colored as in Figure 3c) of one larva during one stimulation trial comprising 5 photostimulation events (red vertical bars at the bottom of the map). Warmer colors indicate higher activity. **b.** Chord diagram representing all-against-all correlations of neuronal activity of brain regions (nodes). Link color and width represents connectivity strength. Self-correlations are excluded from the diagram. Data are presented as mean values of N = 6 ReaChR^+^ larvae. **c.** Seed-based functional connectivity map showing the Pearson’s correlation coefficient of the activity of each brain voxel with average activity extracted from the stimulation site (LHb, seed). Maps are maximum intensity projections of the average 3D map (obtained from individual functional connectivity maps of N = 6 ReaChR^+^larvae, Supplementary Movie 5) along the transverse (left), sagittal (right) and coronal (bottom) axes. Scale bar, 100 μm. **d.** Average Pearson’s correlation coefficient of different brain regions of the larval brain. Correlation values are obtained from individual average functional connectivity maps. Seed of correlation is represented by the average activity extracted from the LHb (stimulation site). Regions are color-coded as in Figure 3c. Gray horizontal bars represent intragroup mean values. *** *P* < 0.0001, * *P* < 0.0345, one way ANOVA and post-hoc Tukey’s test. **e.** Average normalized distributions of Pearson’s correlation coefficient values extracted from voxels inside LHb, IPN and RHb. *** *P* < 0.0001, KS test with Bonferroni correction (α = 0.01667). **f.** Binarized functional connectivity map (Pearson’s seed-correlation coefficient > 0.12 threshold value, separating regions showing significantly higher correlation) of the left habenula (magenta) overlaid on neurons (green) and larval head structure (gray). A, anterior; V, ventral; L, left. N = 6 ReaChR^+^larvae.

Next, we investigated the seed-based functional connectivity of the left habenular nucleus. To this end, we computed Pearson’s correlation between the average neuronal activity of the LHb (seed) and the activity of each brain voxel. Figure 4c shows different projections of the average functional connectivity map of the LHb (Supplementary Movie 5). Besides LHb neurons showing expected high self-correlation, IPN neurons show visible higher functional connectivity with respect to other brain regions. This result is confirmed by the analysis of the average correlation coefficient of the different regions (Figure 3d), where IPN is the only region presenting a statistically significant functional connectivity with the LHb. Figure 3e shows the average normalized distributions of correlation coefficients computed from voxels inside LHb, IPN and RHb. With respect to RHb, which has a distribution basically centered at 0, neurons of LHb and IPN show functional connectivity values as high as 100% and 65%, respectively. Based on these results, Figure 4f shows the binarized functional connectivity map of the left habenular nucleus in larval zebrafish (Supplementary Movie 6).

## Discussion

Dissecting brain functional connectivity requires advanced technology for “reading” and “writing” neuronal activity. Here, we have presented the application of an all-optical 2P system intended for simultaneous imaging and optogenetic control of whole-brain neuronal activity in zebrafish larvae. Our method employs light-sheet microscopy to perform functional imaging, ensuring comprehensive mapping of the entire brain at a significantly improved temporal resolution compared to conventional 2P point-scanning imaging techniques. To elicit precise photoactivation within the larval brain, our light targeting unit utilizes two pairs of AODs enabling the displacement of the focal volume to arbitrary locations. Admittedly, the utilization of AODs for optogenetics has been so far restricted to 1P photostimulation in 2D^48-50^, owing to the drop in transmission efficiency along the optical axis that hinders a homogeneous 2P volumetric excitation. However, as we demonstrated in a previous work^35^, by properly tuning the train of chirped radio frequency (RF) signals which drive the AODs, it is feasible to enhance the uniformity of energy delivery when shifting the focus of the excitation beam. This enhancement has allowed us to proficiently execute optogenetic stimulation of specific targets over a volumetric range of 100×100×100 μm^3^. Notably, an intriguing aspect of our approach is that, owing to the use of remote focusing of the detection objective, the localization of the photostimulation volume remains entirely independent of the sequential acquisition of different brain planes, thus affording greater flexibility in our experimental investigations.

As previously mentioned, our setup exploits 2P excitation both for imaging and optogenetic stimulation. On the imaging side, the use of NIR light to produce the sheet of light leads to a significant reduction of common striping artifacts^51^ that otherwise could severely hinder the interpretation of functional data. On the photostimulation side, the use of nonlinear interaction between light and matter enables precise optical confinement of the stimulation volume, without resorting to narrower genetic control of opsin expression, which is typically required when using 1P excitation^52-55^. In addition to these aspects, the exclusive use of NIR light as an excitation source, in contrast to visible lasers, dramatically diminishes unwanted and uncontrolled visual stimulation since these wavelengths are scantily perceived by most vertebrate species, including zebrafish^56,57^. Nevertheless, we observed that 2P light-sheet imaging can elicit a power-dependent increase in the neuronal activity of zebrafish larvae. This effect may be attributed to non-visual sensory perception of the excitation light, thus remarking the significance of maintaining low the overall energy applied to the sample.

To the best of our knowledge, this is the first time that a fully 2P all-optical setup employs light-sheet microscopy for rapid whole-brain imaging and AODs for 3D optogenetic stimulation.

With the aim of establishing an all-optical paradigm for investigating the functional connectivity of the larval zebrafish brain, we considered different pairs of sensor/actuator and eventually opted for the GCaMP6s/ReaChR couple. The green calcium reporter GCaMP6s represents a reliable indicator which has undergone extensive evaluation^58-61^. On the other hand, the actuator ReaChR, in comparison with other red-shifted opsins, has a slow channel closing mechanism which is particularly suitable for both sequential photostimulation approaches and 2P excitation^23^. A crucial aspect in all-optical studies lies in the separation between the excitation spectra of proteins used for stimulating and revealing neuronal activity. Previous research has demonstrated that the slow channel closing of ReaChR makes this opsin more susceptible to crosstalk activation when scanning the 920 nm imaging laser at power levels exceeding 60 mW^41^. However, in our work, we did not observe a significant increase in cross-activation even at power levels as high as 100 mW. This divergence can be attributed to our use of 2P light-sheet instead of 2P point scanning imaging. In digitally scanned 2P LSFM, the use of low numerical aperture excitation objectives (to obtain a stretched axial illumination PSF, continuously scanned to produce the sheet of light) results in lower intensities (and thus lower photon density) in comparison to point-scanning methods, for equal laser powers. It is worth noting that, despite the negligible crosstalk, 2P light-sheet imaging may still lead to subthreshold activation of ReaChR^+^ neurons, potentially resulting in altered network excitability^62^. Previous studies employed 1030 nm pulsed lasers to stimulate ReaChR^41,63^. The results of our work demonstrate the feasibility of photostimulating ReaChR at 1064 nm, a wavelength red-shifted by almost 100 nm compared to the ReaChR 2P absorption peak (975 nm^63^). Furthermore, the use of the 1064 nm wavelength for photostimulation, which is red shifted with respect to the tail of the 2P excitation cross-section of GCaMP6s^64^, accounts for the absence of fluorescence artifacts potentially caused by the calcium indicator excitation at the wavelength employed for optogenetic intervention. The characterization of the kinetic features of calcium transients elicited by optogenetic stimulation, which served as a benchmark for identifying the optimal excitation configuration, highlighted an interesting aspect. We observed a decrease in the calcium transient rise time in response to longer stimulation durations. This result may be attributed to the fact that ReaChR has a channel off rate (*τ*-off) of 140 ms^37^, enabling it to integrate photons beyond the duration of a single volume iteration (125 ms). Supporting this hypothesis is the fact that, after two iterations over the stimulation volume (250 ms and on), the rise time remains constant.

As the system allows to accurately identify groups of neurons functionally connected with the stimulated ones, we exploited the setup to explore the efferent functional connectivity of the left habenula. The habenulae are bilateral nuclei located in the diencephalon which are highly conserved among vertebrates and connect brain regions involved in diverse emotional states such as fear and aversion, as well as learning and memory^65^. Like mammals, the habenulae in zebrafish are highly connected hubs receiving afferents from entopeduncular nucleus^66^, hypothalamus, and median raphe^67^, in addition to left-right asymmetric inputs^45,68^. The habenula can be divided into a dorsal (dHb) and a ventral (vHb) portion (equivalent to the mammalian medial^69^ and lateral habenula^70^, respectively), each exhibiting exclusive efferent connections. Specifically, dHb sends inputs to the IPN while vHb projects to the median raphe^69,70^. As a consequence, optogenetic stimulation of the entire LHb should lead, in principle, to the activation of both IPN and raphe. However, in our experiments, we see a high probability of activation and a strong correlation only within the IPN population of neurons. This apparent discrepancy can be explained by the fact that, at the larval stage, the vHb represents only a small fraction of the overall habenular volume^70^. As a result, the limited number of vHb neurons would possess a reduced number of connections with the median raphe, resulting in a weak downstream communication. Furthermore, as described by Amo and colleagues^70^, although vHb neurons terminate in the median raphe, no direct contact with serotonergic neurons is observed, suggesting the presence of interneurons which may bridge the link, similar to what is observed in mammals^71^. This inhibitory connection would be coherent with the absence of activation of the raphe which we observed upon left habenula stimulation. Notably, we did not observe any activation in regions downstream to the IPN either. Although adult zebrafish exhibit IPN habenular-recipient neurons projecting to the dorsal tegmental area (DTA) or griseum centrale (GC)^72^, our results corroborate the structural observations of Ma and colleagues from a functional standpoint. Indeed, using anterograde viral labeling of postsynaptic targets, Ma et al. highlighted that in larval zebrafish habenular-recipient neurons of the IPN do not emanate any efferent axon^46^.

LHb and IPN show a high interindividual variability in terms of average activation probability while a lower variability degree in terms of correlation. This point is due to the fact that larvae may present slightly different opsin expression levels, which result in higher variance in the amplitude of evoked calcium transients and, thus, higher activation probability (i.e., the probability to exceed an arbitrary amplitude threshold). Conversely, the strength of functional connection (i.e., the degree of correlation) appears to be not dependent on the amplitude of evoked neuronal activity. This aspect is also confirmed by the high cross power spectral density in a narrow bandwidth centered on the frequency of the triggered optogenetic stimulus, which we observed in the average activity time traces extracted from LHb and IPN.

In conclusion, we employed optogenetic stimulation to map the whole-brain functional connectivity of the left habenula efferent pathway in zebrafish larvae. This application has showcased the remarkable capabilities of our 2P setup for conducting crosstalk-free all-optical investigations. The use of AODs for precisely addressing the photostimulation is a hot topic in system neuroscience as evidenced by recent conference contributions^73,74^. These devices indeed, owing to their discontinuous scanning with constant access time, enable random-access modality. This feature empowers AODs with the native capability to perform rapid sequential excitation over multiple sparsely distributed cellular targets, a feature recently sought after also by SLM adopters^75^. Indeed, rapid sequential stimulation enabled by AODs represents an invaluable tool for studies aiming at replicating a physiological neuronal activation pattern.

Future efforts will be devoted to push further the volume addressable with AODs scanning, while concurrently improving the uniformity of energy delivery. Furthermore, leveraging transgenic strains that express the actuator under more selective promoters (such as *vglut2* for glutamatergic and *gad1b* for GABAergic neurons) will undoubtedly help producing accurate inferences on network structures^76,77^, thus boosting the quest towards a comprehensive picture of zebrafish brain functional connectivity. Together, light-sheet microscopy and 3D optogenetics with AODs, along with the employment of larval zebrafish, offer a promising avenue for bridging the gap between microscale resolution and macroscale investigations, enabling the mapping of whole-brain functional connectivity at previously unattainable spatio-temporal scales.

## Methods

### Optical setup

The all-optical control and readout of zebrafish neuronal activity is achieved through a custom system that combines a 2P dual-sided illumination LSFM for whole-brain calcium imaging^40,78,79^ and an AOD-based 2P light-targeting system for 3D optogenetic stimulation^35^ (Figure 1a). The two systems have been slightly modified with respect to the previous published versions to optically coupling them. Briefly, the 2P light-sheet imaging path is equipped with a pulsed Ti:Sa laser (Chameleon Ultra II, Coherent), tuned at 920 nm. After a group delay dispersion precompensation step, the near-infrared beam is adjusted in power and routed to an electro-optical modulator (EOM) employed to switch the light polarization orientation between two orthogonal states at a frequency of 100 kHz. A half-wave plate and a quarter-wave plate are used to control the light polarization plane and to pre-compensate for polarization distortions. Then, the beam is routed to a hybrid pair of galvanometric mirrors (GMs). One is a fast resonant mirror (CRS-8 kHz, Cambridge Technology) used to digitally generate the virtual light-sheet scanning (frequency 8 kHz) the larva along the rostro-caudal direction. The second GM is a closed-loop mirror (6215H, Cambridge Technology) used to displace the light-sheet along the dorso-ventral direction. The scanned beam is driven by a scan lens and a tube lens into a polarizing beam splitter which diverts the light alternatively into either of the two excitation arms, according to the instantaneous polarization state imposed by the EOM. In order to maximize fluorescence collection, after the beam splitter in one of the two arms a half-wave plate is used to rotate the light polarization plane so that light coming from both the excitation paths is polarized parallel to the table surface^39^. Through a twin relay system, the beams are finally routed into the excitation objectives (XLFLUOR4X/340/0,28, Olympus). The excitation light is focused inside a fish water-filled custom imaging chamber, thermostated at 28.5 °C. The fine positioning of the sample under the detection objective is performed with three motorized stages. Fluorescence emitted by the sample is collected with a water-immersion objective (XLUMPLFLN20XW, Olympus, NA = 1). Finally, a relay system brings the collected signal to an electrically tunable lens (ETL; EL- 16-40-TC-VIS-5D-C, Optotune) which performs remote axial scanning of the detection objective focal plane in sync with the light-sheet closed-loop displacement. The signal collected is filtered (FF01-510/84-25 nm BrightLine® single-band bandpass filter, Semrock) to select green emission. Filtered light reaches an air objective (UPLFLN10×2, Olympus, NA = 0.3) which demagnifies the image onto a subarray (512×512 pixels) of an sCMOS camera (ORCA-Flash4.0 V3, Hamamatsu) working at 16-bit depth of integer gray levels. The final magnification of the imaging system is 3×, with a resulting pixel size of 2.2 μm.

The 3D light-targeting system employs a 1064 nm pulsed laser (FP-1060-5-fs Fianium FemtoPower, NKT Photonics, Birkerød, Denmark) as an excitation source. The output power (max. 5W) is attenuated and conveyed to a half-wave plate, employed to adjust the polarization of the beam, before entering the first AODs stage (DTSXY-400 AA Opto Electronic, Orsay, France). The output beam is then coupled with the second AODs stage through two 1:1 relay systems. From the exit of the second stage, by means of a 1:1 relay system, the beam is routed to a couple of galvanometric mirrors (GVS112, Thorlabs). The scanned beam is then optically coupled with a scan lens (AC254- 100-B, Thorlabs) and a tube lens (F = 300 mm, in turn formed by two achromatic doublets - AC254- 150-C-MLE, F = 150 mm, Thorlabs so customized to avoid aberrations). The excitation light is finally deflected by a dichroic mirror (DMSP926B, Thorlabs) toward the back pupil of the illumination objective, which is also employed by the imaging system for fluorescence detection.

### Optical characterization of the system

A detailed optical characterization of the 2P light-sheet system was described in a previous work of our group^40^. Herein, we describe the optical performance of the AOD-based light-targeting system used for optogenetic stimulation. When using AODs to move the beam away from its native focus, the illumination axial displacement - or defocus - has a linear relation with the chirp parameter α, i.e. the rate of frequency change of the driving radio waves^80^. We thus measured the axial displacement of the focused beam as a function of α, by illuminating a fluorescent solution (Sulforhodamine 101; S7635, Sigma-Aldrich) and localizing the maximum fluorescent peak in the volume as a function of α, spanning from ™1 MHz/µs to 1 MHz/µs (step size 0.1 MHz/µs). For each chirp configuration, the ETL in detection was used to obtain a 200-µm deep stack (step size: 1 µm) centered at the nominal focal plane of the illumination objective. Supplementary Figure 5a shows the axial position of the fluorescent intensity peak as a function of the chirp addressed, following an expected linear trend. We evaluated the conversion coefficient from the slope of the linear fit, which was 50.44 ± 3.45 µm/MHz/µs (mean ± sd).

We also measured the amount of energy released on the sample as a function of the imposed chirp, or basically, as a function of the time spent illuminating axially displaced targets. Indeed, the beam would spend slightly different periods lighting spots displaced in different *z*-planes as the effective ramp time is a function of the chirp α imposed on the driving RF signal. As explained in detail in a previous work^35^, we partially recovered this non-uniformity in the distribution of power deposited along the axial direction by repeatedly triggering equal frequency ramps within the desired dwell time (here 20 µs each point), using what we called multi trigger modality. In such a way, with respect to conventional single trigger modality, we effectively multiplied the minimum energy deposited on different focal planes, while keeping a stable dwell time. Supplementary Figure 5b shows in black the usual light transmission distribution collected as a function of the chirp (single trigger modality) and in blue the distribution obtained with our multi-trigger approach.

We then measured the point spread function (PSF) of the light-targeting system using subdiffraction-sized fluorescent beads (TetraSpeck microspheres, radius 50 nm; T7279, Invitrogen) embedded in agarose gel (1.5 %, w/v) at a final concentration of 0.0025% (vol/vol). The measure was performed on a field of view of 100×100 μm^2^, performing raster scans of 500×500 points. The objective was moved axially covering a 200 μm range (*z*-step: 1 μm) and the emitted signal was conveyed and collected on an auxiliary photomultiplier tube positioned downstream to the fluorescence-collecting objective. Radial and axial intensity profiles of 25 beads were computed using the open-source software ImageJ^81^ and fitted with Gaussian functions in Origin Pro 2021 (OriginLab Corp.) to estimate the full width at half maximum (FWHM). Supplementary Figure 5c shows, as an example, raw fluorescence distributions of 5 beads and the Gaussian fit corresponding to the average FWHM, plotted in red and black for the radial and axial PSF, respectively. We found them to be FWHMr = 0.81 ± 0.06 µm and FWHMa = 3.79 ± 0.66 µm (mean ± sd). This measurement was performed by setting static frequencies on the AOD driving RF signals. In order to evaluate eventual illumination spatial distortions arising away from the objective nominal focal plane, we repeated the same PSF measurement for different chirps, or in other terms, for different AOD-controlled axial displacements (80 µm range, step size 20 µm). The average FWHM obtained on the bead intensity distribution is shown in Supplementary Figure 5d. The radial PSF of the system remains approximately constant as a function of the chirp parameter. A small change is due to the chromatic dispersion affecting the laser beam interacting with the crystal. The deflection angle induced by the AODs to the incident beam is frequency and wavelength dependent. This means that a broadband laser is straightforwardly spatially dispersed by the crystal and that the frequency variations can slightly affect this distortion. Moreover, the axial PSF tends to become slightly oblong at increasing axial displacement. This effect is attributable to the temporal dispersion affecting a short-pulsed laser beam interacting with the crystal. This temporal broadening reduces the axial 2P excitation efficiency generating a larger axial PSF. This effect is more evident when a chirp is applied to the RF signals driving the AODs. In this condition the beam reaches the objective back-pupil in a non-collimated state. Future efforts will be devoted to the compensation of chromatic aberration and temporal dispersion, for example employing a highly dispersive prism upstream to AODs^82^.

### Zebrafish lines and maintenance

The double *Tg(elavl3:H2B-GCaMP6s; elavl3:ReaChR-TagRFP)* zebrafish line was obtained from outcrossing the *Tg(elavl3:H2B-GCaMP6s)^20,83^*and the *Tg(elavl3:ReaChR-TagRFP)^35,58^* lines on slc45a2^b4/-^ heterozygous *albino* background^84^, which we previously generated. The double transgenic line expresses in all differentiated neurons the fluorescent calcium reporter GCaMP6s (nucleus) and the red-shifted light-activable cation channel ReaChR (plasma membrane). ReaChR is expressed as a fusion peptide with red fluorescent protein TagRFP to make it localizable. Zebrafish strains were reared according to standard procedures^85^, fed twice a day with dry food and brine shrimps nauplii (*Artemia salina*), both for nutrition and environmental enrichment. For experiments we employed n = 16, 5 dpf *Tg(elavl3:H2B-GCaMP6s; elavl3:ReaChR-TagRFP)* and n = 9, 5 dpf *Tg(elavl3:H2B-GCaMP6s)*, both on slc45a2^b4/b4^ homozygous *albino* background. Zebrafish larvae used in the experiments were maintained at 28.5 °C in fish water (150 mg/L Instant Ocean, 6.9 mg/L NaH2PO4, 12.5 mg/L Na2HPO4, 1 mg/L methylene blue; conductivity 300 μS/cm, pH 7.2) under 14/10 light/dark cycle, according to standard protocols^85^. Experiments involving zebrafish larvae were carried out in compliance with European and Italian laws on animal experimentation (Directive 2010/63/EU and D.L. 4 March 2014, n.26, respectively), under authorization n.606/2020- PR from the Italian Ministry of Health.

### Zebrafish larvae preparation

In order to select calcium reporter/opsin-expressing larvae to be used in the experiments, 3 dpf embryos were subjected to a fluorescence screening. Embryos were first slightly anesthetized with a bath in tricaine (160 mg/L in fish water; A5040, Sigma-Aldrich) to reduce movements. Using a stereomicroscope (Stemi 508, Carl Zeiss) equipped with LEDs for fluorescence excitation (for GCaMP6s: blue LED, M470L3; for TagRFP: green LED, M565L3, both from Thorlabs) and fluorescence filters to block excitation light (for GCaMP6s: FF01-510/84-25; for TagRFP: FF01- 593/LP-25, both from Semrock), embryos were selected according to the presence of brighter green/red fluorescent signals in the central nervous system. Screened embryos were transferred to a Petri dish containing fresh fish water and kept in an incubator at 28.5 °C until 5 dpf. Zebrafish larvae were mounted as previously described^86^. Briefly, each larva was transferred into a reaction tube containing 1.5% (w/v) low-gelling temperature agarose (A9414, Sigma-Aldrich) in fish water, maintained fluid on a heater set at 38°C. Using a plastic pipette, larvae were then placed on a microscope slide inside a drop of melted agarose. Before gel polymerization, their position was adjusted using a couple of fine pipette tips in order for the dorsal portion to face upwards. To avoid movement artifacts during measurements, larvae were paralyzed by a 10-minute treatment with 2 mM d-tubocurarine (93750, Sigma-Aldrich), a neuromuscular blocker. The mounted larva was then placed inside the imaging chamber filled with fish water, thermostated at 28.5 °C for the entire duration of the experiment.

### Simultaneous calcium imaging and optogenetic stimulation

Whole-brain calcium imaging was performed at 2.5 Hz with 41 stacked z-planes spanning a depth of 200 μm. An interslice spacing of 5 μm was chosen as it coincides with the half width at half maximum of the detection axial PSF. Before each measurement, the scanning amplitude of the resonant galvo mirror was tuned in order to produce a virtual light-sheet having a length matching the size of the larval brain in the rostro-caudal direction. The laser wavelength was set to 920 nm to optimally excite GCaMP6s fluorescence. Unless otherwise stated, power of the 920 nm laser was set to 60 mW on the sample.

Optogenetic stimulation was performed at 1064 nm with a laser power at the sample of 30 mW (unless otherwise specified). Before each experimental session, the 1064 nm stimulation laser was finely aligned to be at the center of the camera field-of-view. Then, by means of the galvo mirrors present in the stimulation path, the offset position of the stimulation beam was coarsely displaced in the *x*-*y* direction towards the center of the area to be stimulated. During the optogenetics experiment the stimulation volume was covered by discontinuously scanning the beam focus via the two couples of AODs. A typical volume of 50×50×50 μm^3^ was covered with 6250 points (point x-y density: 1 point/0.25 μm^2^; z-step: 5 μm) with a point dwell time of 20 μs (overall time: 125 ms). The medial plane of the stimulation volume (chirp = 0 MHz/μs, null defocus) was adjusted to overlap with the medial plane of the LHb. Unless otherwise stated, each stimulus consisted of 4 complete cycles of the entire volume, lasting 500 ms. Each stimulation trial consisted of 100 s of whole-brain calcium imaging during which 5 optogenetic stimuli (interstimulus interval: 16 s) were performed in the same volumetric site. Six trials were performed on each larva, with an intertrial interval ranging from 1 to 3 min. Overall, each larva was imaged for 10 min during which it received 30 stimuli.

### Structural imaging to evaluate expression patterns in double transgenic zebrafish larvae

Confocal imaging of a 5 dpf *Tg(elavl3:H2B-GCaMP6s; elavl3:ReaChR-TagRFP)* larva on albino background was performed in order to evaluate spatial expression of the two proteins. The larva was mounted in agarose as described above and deeply anesthetized with tricaine (300 mg/L in fish water). We employed a commercial confocal microscope (T*i*2, Nikon) equipped with two continuous wavelength lasers emitting at 488 and 561 nm for GCaMP6s and TagRFP excitation, respectively. Imaging was performed using a 10× objective allowing the entire head of the animal to fit into the field-of-view. Using a piezo-electric motor (PIFOC, Physik Instrumente - PI) the objective was moved at 182 consecutive positions (z-step: 2 μm) in order to acquire the volume of the larval head.

### Data analysis

#### Preprocessing

Whole-brain calcium imaging data were processed as follows. Images composing the hyperstacks were first 2×2 binned (method: average) in the *x* and *y* dimensions to obtain quasi-isotropic voxel size (4.4×4.4×5 μm^3^). Then, employing a custom tool written in Python 3, we computed the voxel-wise *ΔF/F0* of each volumetric recording, after background subtraction. *F0* was calculated using FastChrom’s baseline estimation method^87^.

#### Quantification of imaging crosstalk and optogenetic activation extent/specificity

To quantify crosstalk during imaging we first computed the standard deviation (SD) of each voxel composing the larval brain along 5-min whole-brain calcium imaging. The distribution of SD values of each brain was first normalized with respect to the total number of voxels and then pooled (method: average) according to the larval strain (ReaChR^+^ and ReaChR^−^). Similarly, the normalized distributions of SD values for R^+^ and R^−^ larvae subjected to 100-s whole-brain imaging during which they received 5 photostimulations (1064 nm) were calculated to evaluate the effect of the optogenetic stimulation. Imaging crosstalk and optogenetic stimulation indices were calculated using the Hellinger distance equation^88^ as a measure of the distance between two probability distributions P and Q:

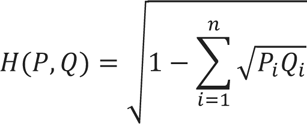

Errors of the Hellinger distances were calculated according to error propagation theory as follows:

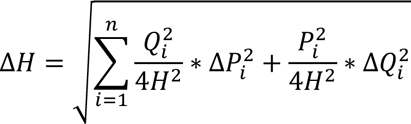

Finally, normalized distributions of SD values for R^−^ larvae exposed either to imaging (100 s) only or to imaging and photostimulation (100 s and 5 stimuli at 1064 nm) were calculated to evaluate the specificity of the effect observed.

#### Characterization of stimulation-induced calcium transients

To characterize the neuronal activation as a function of the stimulation parameters (scan time and laser power), we first extracted the fluorescence time traces averaged over the entire stimulation site (i.e., left habenula) from 4D *ΔF/F0* hyperstacks. Time traces were windowed for isolating and aligning the 3 stimulation events contained into a single trial. Isolated calcium transients were analyzed using the *peak analyzer* function in Origin Pro 2021 (OriginLab Corp.) to obtain peak amplitude, rise/decay time (i.e., time from baseline to peak and time from peak to baseline, respectively) and duration values. Pooled peak duration data were obtained by first averaging 3 events of the same larva (intra-individual) and then averaging between larvae (inter-individual).

#### Activation probability and correlation maps

Using a custom Python tool, we calculated the probability of each voxel composing the brain to be active in response to the optogenetic stimulation. For each stimulation event, a voxel was considered active if its change in fluorescence in a 2 s time window after the stimulation exceeded 3 standard deviations above its baseline level (2 s pre-stimulation). Only events in which the voxels inside the stimulation volume met the activation criterion were considered effective optogenetic stimulations. By iterating this process for all the stimulation events performed (on the same site of the same larva), we thus calculated the activation probability of each voxel as the number of times the voxel exceeded the threshold divided by the total number of valid stimulations. Employing a second Python tool, we then computed activity correlation maps showing the Pearson’s correlation coefficient between each voxel and the activity extracted from the stimulation site (seed). The 3D maps of correlation and activation probability obtained were then aligned. First, the acquired 4D hyperstacks were time averaged. Secondly, the resulting 3D stack of each larva was registered to a reference brain. Nonrigid image registration was performed using the open source software Computational Morphometry Toolkit (CMTK 3.3.1, https://www.nitrc.org/projects/cmtk/) and the ImageJ user interface^89^, employing the command string (-awr 01 −X 52 −C 8 −G 80 −R 3 −A ’--accuracy 1.6’ -W ’-- accuracy 0.4’). Calculated morphing transformations were finally applied to the corresponding 3D maps.

Following the zebrafish brain atlases^90,91^, the volumetric regions of interest (ROIs) used in the analysis were manually drawn onto the reference brain (employing ImageJ), based on anatomical boundaries. The 10 volumetric ROIs were then adopted to extract from each map the pixel-wise distribution of activation probability/correlation coefficient values used for further analyses.

The binarized functional connectivity map shown in Figure 4f was obtained after applying a threshold on the Pearson’s correlation coefficient to the average correlation map shown in Figure 4c. The 0.12 value represented the correlation coefficient threshold separating regions showing significantly higher connectivity.

#### Wavelet cross-spectrum analysis

The possible coupling between the delineated brain ROIs and the stimulation site was characterized also in the spectral domain by quantifying and inspecting their wavelet cross power spectral density^92^. The wavelet transforms of the average activity signals extracted from each ROI were computed using the Morlet mother wavelet, adopting a central frequency f0 = 1 Hz as time-frequency resolution parameter, and 256 voices per octave for a fine frequency discretization. Spurious time-boundary effects were tackled by first applying a zero-padding scheme to the original time series, and then isolating the so-called cone of influence, i.e. the time–frequency region where boundary distortions in the cross power spectral density estimates are negligible^93^.

#### Statistics and Reproducibility

To guarantee reproducibility of the findings and avoid bias, larvae employed in the experiments never belonged to a single batch of eggs. Expression pattern of GCaMP6s and ReaChR was evaluated on N = 1 ReaChR^+^ larva by confocal imaging. Crosstalk activation of ReaChR by the 920 nm excitation light-sheet imaging was evaluated on N = 3 ReaChR^+^ and N = 3 ReaChR^−^ larvae. Effect of the optogenetic stimulation was evaluated on N = 6 ReaChR^+^ and N = 6 ReaChR^−^ larvae. Characterization of optogenetically induced calcium transients as a function of stimulation settings was performed on N = 4 ReaChR^+^ larvae (n = 3 calcium transients per larva). Activation probability and correlation were evaluated on N = 6 ReaChR^+^ (n = 30 stimulations per larva).

OriginPro 2021 (OriginLab Corp.) was used to carry out all statistical analyses. Unless otherwise stated, results were considered statistically significant if their corresponding *p*-value was less than 0.05 (* P < 0.05; ** P < 0.01; *** P < 0.0001). Both intergroup and intragroup statistical significance of imaging crosstalk (Figure 1e) were performed using two-way ANOVA (factors: zebrafish strain, imaging power) followed by post-hoc comparisons with Tukey’s method. For intergroup statistical evaluations of both activation probability (Figure 3c) and Pearson’s correlation coefficient (Figure 4d), we first verified normality distribution of data using the Shapiro-Wilk test (see Supplementary Figure 6a,b for test results) and then performed one way ANOVA (factor: brain region), followed by post-hoc comparisons employing Tukey’s method. Statistical comparisons of median SD values to evaluate effect of the optogenetic stimulation (Figures 1f-g and S2) were performed using unpaired *t*-test.

Statistical comparisons of SD (Figure 3d) and Pearson’s correlation coefficient (Figure 4e) average distributions were performed with the two-sample Kolmogorov-Smirnov test (KS test), applying Bonferroni correction (α = 0.05/3 = 0.01667, in both cases).

## Data availability

Data underlying plots, custom-written LabVIEW/Python/C++ codes and all other relevant data are available from the corresponding authors upon reasonable request.

## Notes

The authors declare no conflicts of interest.

## Author contributions

LT and FSP conceived the study, LT designed the experiments, LT performed the experiments, LT and PR performed characterization of calcium activity response to optogenetic stimulation, PR optically coupled the stimulation and imaging systems, PR performed optical characterization of the photostimulation system, MS developed the Python and C++ software of the custom RF signal generator driving the AOD-based photostimulation system, MS and MM programmed the LabVIEW software used for operating the photostimulation system, LT conceived data analysis, MS developed the Python tools for *ΔF/F0* calculation, correlation and wavelet cross-spectrum analyses, GdV programmed the Python software for activation probability computation, LT and GdV set up statistical analysis, LT, MS and PR performed data analysis, FV supervised zebrafish experiments, FSP acquired funding, LT wrote the manuscript. All authors reviewed and agreed to the published version of the manuscript.

## Acknowledgements

We thank Dr. Ludovico Silvestri (University of Florence), Dr. Leonardo Sacconi (INO-CNR), and Domenico Alfieri (Light4Tech) for fruitful discussions regarding AODs implementation. We thank Dorotea Nardini (University of Florence) for the helpful discussion about error propagation theory. This project was funded by: the European Research Council (ERC) under the European Union’s Horizon 2020 Research and Innovation program [Grants n.692943 (BrainBIT) and n.966623 (DAPTOMIC)]; the Italian Ministry for Education, University, and Research in the framework of the Advanced Light Microscopy Italian Node of Euro-Bioimaging ERIC; Bank Foundation Fondazione Cassa di Risparmio di Firenze with grants “Human Brain Optical Mapping”.

## Supplementary Movies legends

**Supplementary Movie 1. Expression patterns of the double transgenic *Tg(elavl3:H2B-GCaMP6; elavl3:ReaChR-TagRFP) zebrafish line*.**

Confocal stack (z-step: 2 μm) of the head of a 5 dpf *Tg(elavl3:H2B-GCaMP6; elavl3:ReaChR-TagRFP)* zebrafish larva. In green, the nuclear localized fluorescent calcium indicator GCaMP6s. In magenta, the membrane light-gated cation channel ReaChR (expressed as a fusion protein with TagRFP). The *elavl3* promoter drives the expression of both the indicator and the opsin in all differentiated neurons. Scale bar: 100 μm.

**Supplementary Movie 2. Raw timelapse acquisition of whole-brain imaging during optogenetic stimulation of the left habenula.**

Composite showing the raw timelapse of 8 evenly distributed *z*-planes of the larval zebrafish brain during a photostimulation trial. Plane depth reported in orange is expressed in μm from the surface of the brain. Stimulation events are indicated by the red dot on the right. Arrowheads indicate the stimulation site (left habenula, blue) and the deep midbrain interpeduncular nucleus (IPN, yellow), described in further analyses. Movie is played at 10× speed. Scale bar, 200 μm.

**Supplementary Movie 3. 3D rotation of whole-brain activation probability map in response to left habenula stimulation.**

The rotating map shows the average probability of each voxel composing the larval brain to be active as a consequence of optogenetic stimulation of the left habenula. Probability is color mapped as in the color bar (black = 0, white = 1). Map rotates along the larval antero-posterior axis. N = 6 ReaChR^+^ larvae.

**Supplementary Movie 4. Volumetric activation probability maps of individual larvae in response to left habenula stimulation.**

Composite showing volumetric activation probability maps of individual larvae. Probability is color-mapped as in the color bar. Each map was obtained calculating the probability of activation on n = 30 photostimulation events.

**Supplementary Movie 5. Volumetric correlation maps of individual larvae.**

Composite of volumetric maps of individual larvae showing the grade of correlation with the optogenetically-induced activity of the left habenula (seed). Pearson’s correlation coefficient is color-mapped as in the color bar. Each map was obtained calculating the correlation with the seed over n = 30 photostimulation events.

**Supplementary Movie 6. 3D rotation of whole-brain functional connectivity of the left habenula.**

The rotating map shows the binarized functional connectivity map of the left habenular nucleus (magenta) overlaid to the brain structure (gray). The map was obtained by binarizing the average seed-based correlation map shown in Figure 4c (Pearson’s correlation coefficient threshold: 0.12). Map rotates along the larval antero-posterior axis. N = 6 ReaChR^+^ larvae.

## Supplementary Figures

**Supplementary Figure 1.**
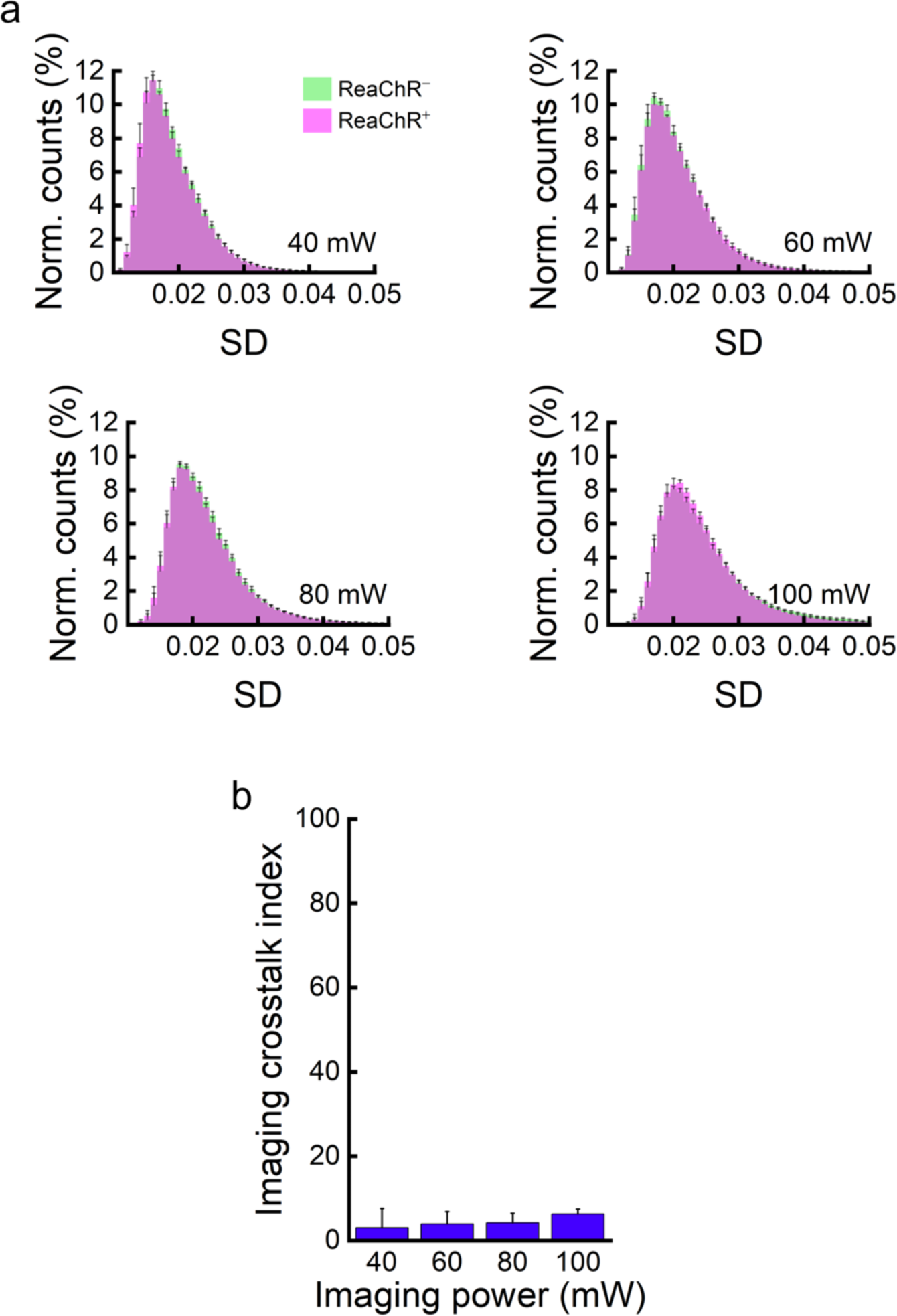
Neuronal activity levels at increasing imaging power. **a.** Average normalized distributions of the SD values of larval brain voxels over 5 minutes of continuous light-sheet imaging (volumetric rate: 2.5 Hz, λex: 920 nm) at laser powers of 40, 60, 80 and 100 mW for ReaChR^+^ and ReaChR^−^ larvae. N = 3 ReaChR^+^ and 3 ReaChR^−^ larvae. **b.** Imaging crosstalk index (Hellinger distance between ReaChR^+^ and ReaChR^−^ distributions) computed for the different power levels tested. Error bars were calculated according to uncertainty propagation theory (see Methods for details).

**Supplementary Figure 2.**
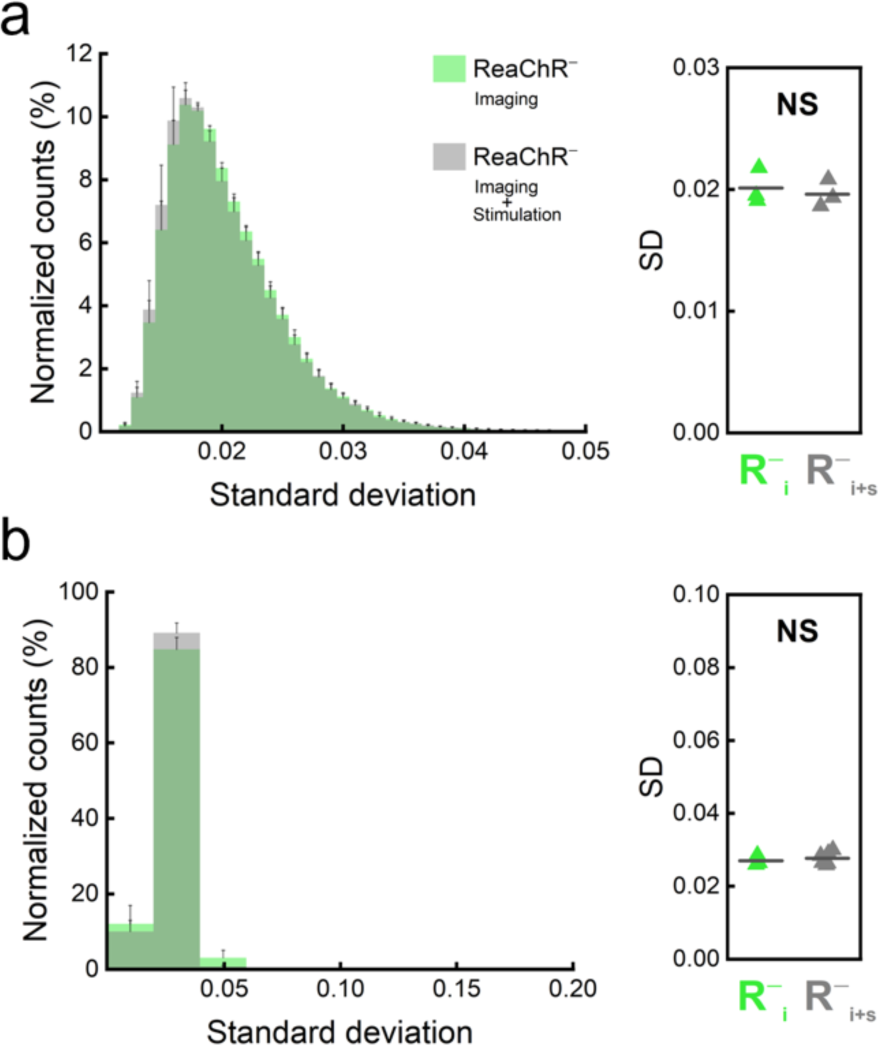
Specificity of the optogenetic stimulation. **a. Left,** average normalized distributions of the SD values of voxels composing the brain of ReaChR^−^ larvae exposed to imaging only (green) or to imaging and photostimulation (gray). **Right,** comparison of median SD values calculated from individual brain-wide SD distributions of ReaChR^−^ larvae during imaging (R^−^i) or imaging and stimulation (R^−^i+s). Intergroup comparisons highlight no statistically significant difference. NS *P* = 0.6510, unpaired *t* test. **b. Left,** average normalized distributions of SD values inside the stimulation site of ReaChR^−^ larvae exposed to imaging only (green) or to imaging and photostimulation (gray). **Right,** comparison of median SD values calculated from individual SD distributions at the stimulation site of ReaChR^−^ larvae undergoing imaging (R^−^i) or both imaging and stimulation (R^−^i+s). Intergroup comparisons highlight no statistical significance. NS *P* = 0.5584, unpaired *t* test.

**Supplementary Figure 3.**
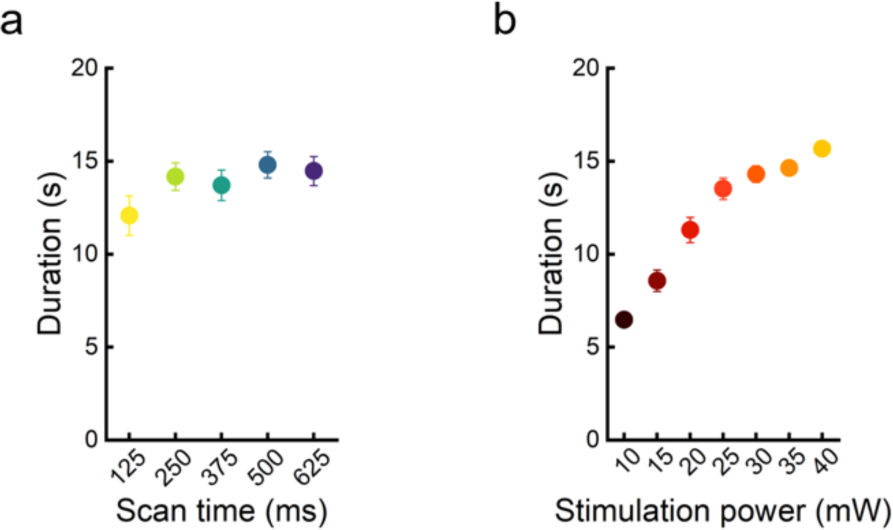
Duration of calcium transients evoked by photostimulation. Average duration of calcium transients as a function of scan time (**a**) and stimulation power (**b**). Symbols are color coded as in Figure 2. Data are presented as mean ± sem. Where not visible, error bars are smaller than symbol size. N = 4 ReaChR^+^ larvae, n = 3 calcium transients per larva.

**Supplementary Figure 4.**
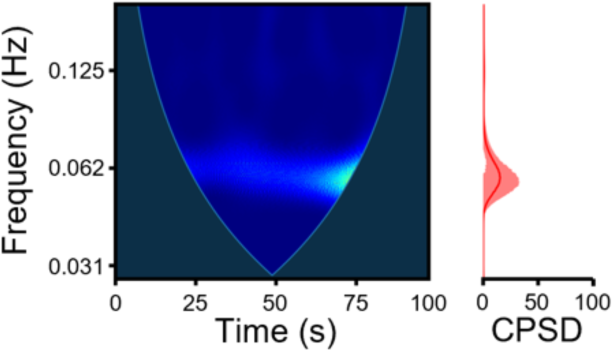
Cross power spectrum of left and right habenula activity. **Left,** map of the cross power spectrum of the LHb and RHb time traces. Warmer colors indicate higher spectral density. **Right,** cross power spectral density (CPSD) averaged over time of the cross-spectrum map. No appreciable cross spectral density is observed, meaning that the two traces do not share significant frequency components. N = 1 ReaChR^+^ larva.

**Supplementary Figure 5.**
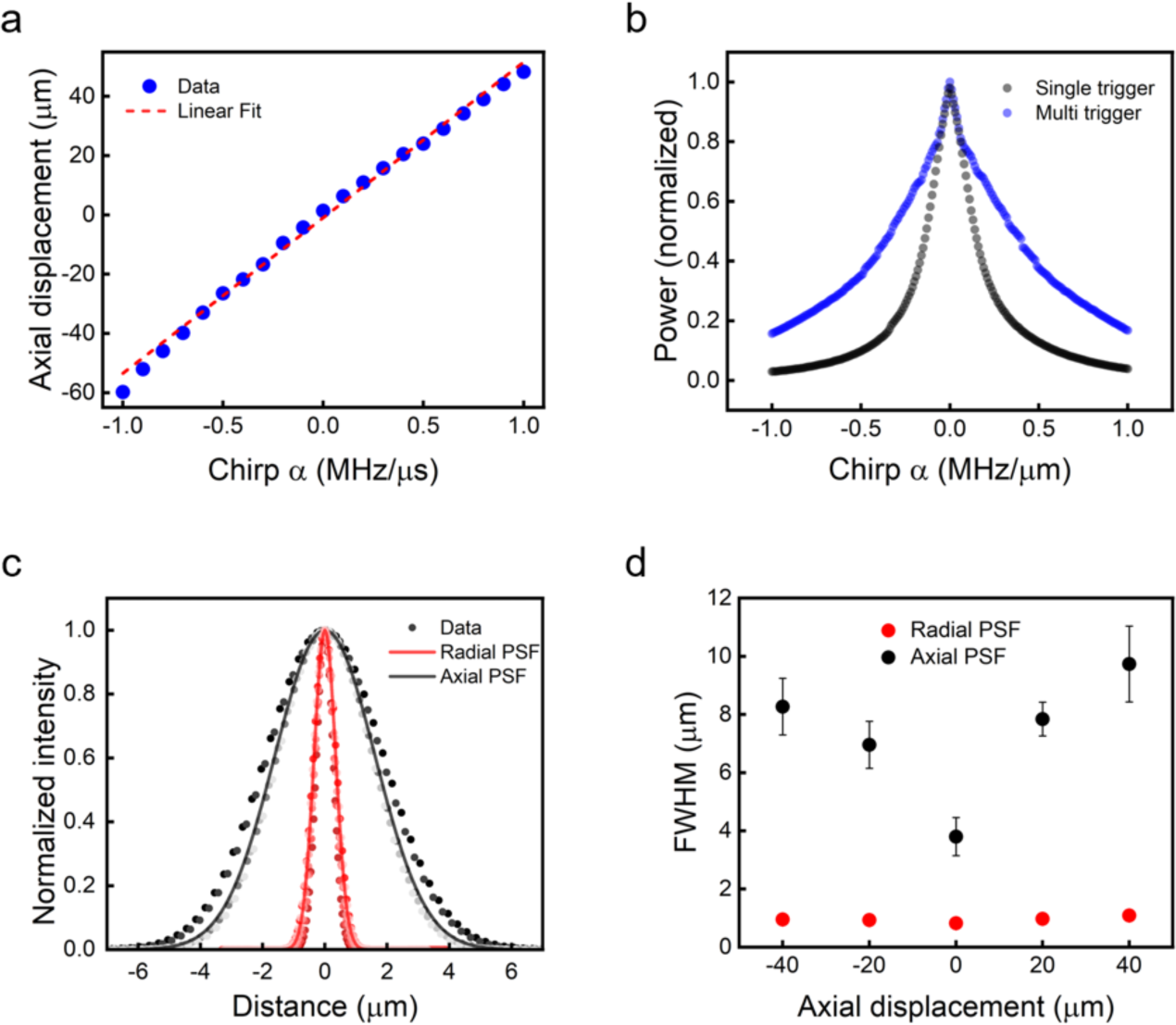
Optical characterization of the light-targeting path. **a.** Axial displacement of the focal point as a function of the chirping frequency applied to AODs. R^2^ = 0.99385- Data are presented as mean ± sem. **b.** Normalized laser power as a function of chirp frequency applied. The ™1/1 MHz/μs chirp range corresponds in our system to a defocusing range of 100 μm. **c.** System point spread function (PSF) is represented as the normalized intensity of subdiffraction-sized fluorescent beads as a function of distance from the center of the bead. Results from 5 beads are displayed. Gaussian fits are calculated on N = 25 fluorescent beads. **d.** FWHM of the PSF as a function of axial displacement (80 μm range) with respect to native focal plane. Data are presented as mean ± sem. N = 25 fluorescent beads. Where not visible, error bars are smaller than symbol size.

**Supplementary Figure 6.**
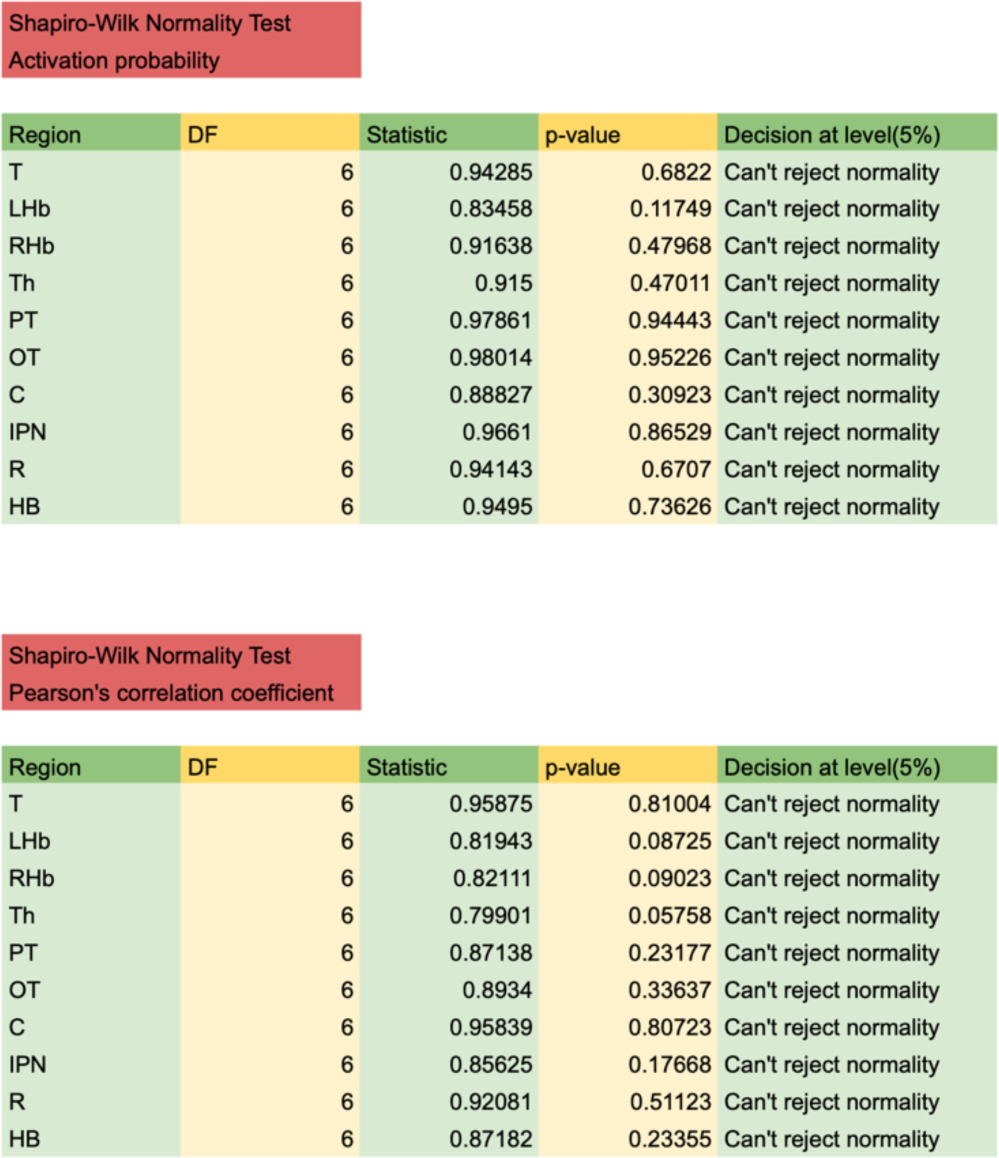
Normality tests. Shapiro-Wilk normality tests for activation probability (**a**) and Pearson’s correlation coefficient (**b**) values in different brain regions of N = 6 ReaChR^+^ larvae. Results of the tests in both cases accept the null hypothesis that data were drawn from a normally distributed population.

## References

1 Rizzolatti, G., Fadiga, L., Gallese, V. & Fogassi, L. Premotor cortex and the recognition of motor actions. Brain Res Cogn Brain Res 3, 131–141, doi:10.1016/0926-6410(95)00038-0 (1996).

2 Sargolini, F. et al. Conjunctive representation of position, direction, and velocity in entorhinal cortex. Science 312, 758–762, doi:10.1126/science.1125572 (2006).

3 Fabbri-Destro, M. & Rizzolatti, G. Mirror neurons and mirror systems in monkeys and humans. Physiology (Bethesda) 23, 171–179, doi:10.1152/physiol.00004.2008 (2008).

4 Norman-Haignere, S. V. et al. A neural population selective for song in human auditory cortex. Curr Biol 32, 1470–1484 e1412, doi:10.1016/j.cub.2022.01.069 (2022).

5 Deisseroth, K. Optogenetics: 10 years of microbial opsins in neuroscience. Nat Neurosci 18, 1213–1225, doi:10.1038/nn.4091 (2015).

6 Lin, M. Z. & Schnitzer, M. J. Genetically encoded indicators of neuronal activity. Nat Neurosci 19, 1142–1153, doi:10.1038/nn.4359 (2016).

7 Packer, A. M. et al. Two-photon optogenetics of dendritic spines and neural circuits. Nat Methods 9, 1202–1205, doi:10.1038/nmeth.2249 (2012).

8 Hochbaum, D. R. et al. All-optical electrophysiology in mammalian neurons using engineered microbial rhodopsins. Nat Methods 11, 825–833, doi:10.1038/nmeth.3000 (2014).

9 Lou, S. et al. Genetically Targeted All-Optical Electrophysiology with a Transgenic Cre-Dependent Optopatch Mouse. J Neurosci 36, 11059–11073, doi:10.1523/JNEUROSCI.1582-16.2016 (2016).

10 Fan, L. Z. et al. All-Optical Electrophysiology Reveals the Role of Lateral Inhibition in Sensory Processing in Cortical Layer 1. Cell 180, 521–535 e518, doi:10.1016/j.cell.2020.01.001 (2020).

11 Choi, T. Y., Choi, T. I., Lee, Y. R., Choe, S. K. & Kim, C. H. Zebrafish as an animal model for biomedical research. Exp Mol Med 53, 310–317, doi:10.1038/s12276-021-00571-5 (2021).

12 Power, R. M. & Huisken, J. A guide to light-sheet fluorescence microscopy for multiscale imaging. Nat Methods 14, 360–373, doi:10.1038/nmeth.4224 (2017).

13 Chen, I. W., Papagiakoumou, E. & Emiliani, V. Towards circuit optogenetics. Curr Opin Neurobiol 50, 179–189, doi:10.1016/j.conb.2018.03.008 (2018).

14 Turrini, L., Roschi, L., de Vito, G., Pavone, F. S. & Vanzi, F. Imaging Approaches to Investigate Pathophysiological Mechanisms of Brain Disease in Zebrafish. Int J Mol Sci 24, doi:10.3390/ijms24129833 (2023).

15 Rickgauer, J. P., Deisseroth, K. & Tank, D. W. Simultaneous cellular-resolution optical perturbation and imaging of place cell firing fields. Nat Neurosci 17, 1816–1824, doi:10.1038/nn.3866 (2014).

16 Dal Maschio, M., Donovan, J. C., Helmbrecht, T. O. & Baier, H. Linking Neurons to Network Function and Behavior by Two-Photon Holographic Optogenetics and Volumetric Imaging. Neuron 94, 774–789 e775, doi:10.1016/j.neuron.2017.04.034 (2017).

17 Forster, D., Dal Maschio, M., Laurell, E. & Baier, H. An optogenetic toolbox for unbiased discovery of functionally connected cells in neural circuits. Nat Commun 8, 116, doi:10.1038/s41467-017-00160-z (2017).

18 Jiao, Z. F. et al. All-optical imaging and manipulation of whole-brain neuronal activities in behaving larval zebrafish. Biomed Opt Express 9, 6154–6169, doi:10.1364/BOE.9.006154 (2018).

19 Huisken, J., Swoger, J., Del Bene, F., Wittbrodt, J. & Stelzer, E. H. Optical sectioning deep inside live embryos by selective plane illumination microscopy. Science 305, 1007–1009, doi:10.1126/science.1100035 (2004).

20 Vladimirov, N. et al. Light-sheet functional imaging in fictively behaving zebrafish. Nat Methods 11, 883–884, doi:10.1038/nmeth.3040 (2014).

21 Packer, A. M., Russell, L. E., Dalgleish, H. W. & Hausser, M. Simultaneous all-optical manipulation and recording of neural circuit activity with cellular resolution in vivo. Nat Methods 12, 140–146, doi:10.1038/nmeth.3217 (2015).

22 Shemesh, O. A. et al. Temporally precise single-cell-resolution optogenetics. Nat Neurosci 20, 1796–1806, doi:10.1038/s41593-017-0018-8 (2017).

23 Ronzitti, E. et al. Recent advances in patterned photostimulation for optogenetics. J Optics-Uk 19, doi:ARTN 113001 10.1088/2040-8986/aa8299 (2017).

24 Picot, A. et al. Temperature Rise under Two-Photon Optogenetic Brain Stimulation. Cell Rep 24, 1243–1253 e1245, doi:10.1016/j.celrep.2018.06.119 (2018).

25 Duocastella, M., Surdo, S., Zunino, A., Diaspro, A. & Saggau, P. Acousto-optic systems for advanced microscopy. J Phys-Photonics 3, doi:ARTN 012004 10.1088/2515-7647/abc23c (2021).

26 Katona, G. et al. Fast two-photon in vivo imaging with three-dimensional random-access scanning in large tissue volumes. Nat Methods 9, 201–208, doi:10.1038/nmeth.1851 (2012).

27 Szalay, G. et al. Fast 3D Imaging of Spine, Dendritic, and Neuronal Assemblies in Behaving Animals. Neuron 92, 723–738, doi:10.1016/j.neuron.2016.10.002 (2016).

28 Nadella, K. M. et al. Random-access scanning microscopy for 3D imaging in awake behaving animals. Nat Methods 13, 1001–1004, doi:10.1038/nmeth.4033 (2016).

29 Villette, V. et al. Ultrafast Two-Photon Imaging of a High-Gain Voltage Indicator in Awake Behaving Mice. Cell 179, 1590–1608 e1523, doi:10.1016/j.cell.2019.11.004 (2019).

30 Akemann, W. et al. Fast optical recording of neuronal activity by three-dimensional custom-access serial holography. Nat Methods 19, 100–110, doi:10.1038/s41592-021-01329-7 (2022).

31 Wang, K. et al. Precise spatiotemporal control of optogenetic activation using an acousto-optic device. PLoS One 6, e28468, doi:10.1371/journal.pone.0028468 (2011).

32 Wang, K. et al. Parallel pathways convey olfactory information with opposite polarities in Drosophila. Proc Natl Acad Sci U S A 111, 3164–3169, doi:10.1073/pnas.1317911111 (2014).

33 Hernandez, O., Pietrajtis, K., Mathieu, B. & Dieudonne, S. Optogenetic stimulation of complex spatio-temporal activity patterns by acousto-optic light steering probes cerebellar granular layer integrative properties. Sci Rep 8, 13768, doi:10.1038/s41598-018-32017-w (2018).

34 Conti, E. et al. Combining Optogenetic Stimulation and Motor Training Improves Functional Recovery and Perilesional Cortical Activity. Neurorehabil Neural Repair 36, 107–118, doi:10.1177/15459683211056656 (2022).

35 Ricci, P. et al. Power-effective scanning with AODs for 3D optogenetic applications. Journal of Biophotonics 15, e202100256, doi:10.1002/jbio.202100256 (2022).

36 Chen, T. W. et al. Ultrasensitive fluorescent proteins for imaging neuronal activity. Nature 499, 295–300, doi:10.1038/nature12354 (2013).

37 Lin, J. Y., Knutsen, P. M., Muller, A., Kleinfeld, D. & Tsien, R. Y. ReaChR: a red-shifted variant of channelrhodopsin enables deep transcranial optogenetic excitation. Nat Neurosci 16, 1499–1508, doi:10.1038/nn.3502 (2013).

38 Zipfel, W. R., Williams, R. M. & Webb, W. W. Nonlinear magic: multiphoton microscopy in the biosciences. Nature Biotechnology 21, 1369–1377, doi:10.1038/nbt899 (2003).

39 de Vito, G. et al. Effects of excitation light polarization on fluorescence emission in two-photon light-sheet microscopy. Biomed Opt Express 11, 4651–4665, doi:10.1364/BOE.396388 (2020).

40 de Vito, G. et al. Fast whole-brain imaging of seizures in zebrafish larvae by two-photon light-sheet microscopy. *Biomed. Opt. Express*, BOE 13, 1516–1536, doi:10.1364/BOE.434146 (2022).

41 Chen, I. W. et al. In Vivo Submillisecond Two-Photon Optogenetics with Temporally Focused Patterned Light. J Neurosci 39, 3484–3497, doi:10.1523/JNEUROSCI.1785-18.2018 (2019).

42 deCarvalho, T. N., et al. Neurotransmitter map of the asymmetric dorsal habenular nuclei of zebrafish. Genesis 52, 636–655, doi:10.1002/dvg.22785 (2014).

43 Fore, S. et al. Functional properties of habenular neurons are determined by developmental stage and sequential neurogenesis. Sci Adv 6, doi:10.1126/sciadv.aaz3173 (2020).

44 Bianco, I. H. & Wilson, S. W. The habenular nuclei: a conserved asymmetric relay station in the vertebrate brain. Philos Trans R Soc Lond B Biol Sci 364, 1005–1020, doi:10.1098/rstb.2008.0213 (2009).

45 Dreosti, E., Vendrell Llopis, N., Carl, M., Yaksi, E. & Wilson, S. W. Left-right asymmetry is required for the habenulae to respond to both visual and olfactory stimuli. Curr Biol 24, 440–445, doi:10.1016/j.cub.2014.01.016 (2014).

46 Ma, M., Kler, S. & Pan, Y. A. Structural Neural Connectivity Analysis in Zebrafish With Restricted Anterograde Transneuronal Viral Labeling and Quantitative Brain Mapping. Front Neural Circuits 13, 85, doi:10.3389/fncir.2019.00085 (2019).

47 Buhler, A. & Carl, M. Zebrafish Tools for Deciphering Habenular Network-Linked Mental Disorders. Biomolecules 11, doi:10.3390/biom11020324 (2021).

48 Huang, L. et al. Task Learning Promotes Plasticity of Interneuron Connectivity Maps in the Olfactory Bulb. J Neurosci 36, 8856–8871, doi:10.1523/JNEUROSCI.0794-16.2016 (2016).

49 Allegra Mascaro, A. L., et al. Combined Rehabilitation Promotes the Recovery of Structural and Functional Features of Healthy Neuronal Networks after Stroke. Cell Rep 28, 3474–3485 e3476, doi:10.1016/j.celrep.2019.08.062 (2019).

50 Resta, F. et al. Large-scale all-optical dissection of motor cortex connectivity shows a segregated organization of mouse forelimb representations. Cell Rep 41, 111627, doi:10.1016/j.celrep.2022.111627 (2022).

51 Ricci, P. et al. Removing striping artifacts in light-sheet fluorescence microscopy: a review. Prog Biophys Mol Biol 168, 52–65, doi:10.1016/j.pbiomolbio.2021.07.003 (2022).

52 Zhang, B. B., Yao, Y. Y., Zhang, H. F., Kawakami, K. & Du, J. L. Left Habenula Mediates Light-Preference Behavior in Zebrafish via an Asymmetrical Visual Pathway. Neuron 93, 914–928 e914, doi:10.1016/j.neuron.2017.01.011 (2017).

53 Helmbrecht, T. O., Dal Maschio, M., Donovan, J. C., Koutsouli, S. & Baier, H. Topography of a Visuomotor Transformation. Neuron 100, 1429–1445 e1424, doi:10.1016/j.neuron.2018.10.021 (2018).

54 Andalman, A. S. et al. Neuronal Dynamics Regulating Brain and Behavioral State Transitions. Cell 177, 970–985.e920, doi:10.1016/j.cell.2019.02.037 (2019).

55 Antinucci, P. et al. A calibrated optogenetic toolbox of stable zebrafish opsin lines. Elife 9, doi:10.7554/eLife.54937 (2020).

56 Lewis, P. R. A theoretical interpretation of spectral sensitivity curves at long wavelengths. J Physiol 130, 45–52, doi:10.1113/jphysiol.1955.sp005391 (1955).

57 Jacobs, G. H. The evolution of vertebrate color vision. Advances in Experimental Medicine and Biology 739, 156–172, doi:10.1007/978-1-4614-1704-0_10 (2012).

58 Dunn, T. W. et al. Brain-wide mapping of neural activity controlling zebrafish exploratory locomotion. Elife 5, e12741, doi:10.7554/eLife.12741 (2016).

59 Turrini, L. et al. Optical mapping of neuronal activity during seizures in zebrafish. Sci Rep 7, 3025, doi:10.1038/s41598-017-03087-z (2017).

60 Feierstein, C. E. et al. Dimensionality reduction reveals separate translation and rotation populations in the zebrafish hindbrain. Curr Biol 33, 3911–3925 e3916, doi:10.1016/j.cub.2023.08.037 (2023).

61 Shainer, I. et al. A single-cell resolution gene expression atlas of the larval zebrafish brain. Sci Adv 9, eade9909, doi:10.1126/sciadv.ade9909 (2023).

62 Ronzitti, E. et al. Submillisecond Optogenetic Control of Neuronal Firing with Two-Photon Holographic Photoactivation of Chronos. J Neurosci 37, 10679–10689, doi:10.1523/JNEUROSCI.1246-17.2017 (2017).

63 Chaigneau, E. et al. Two-Photon Holographic Stimulation of ReaChR. Front Cell Neurosci 10, 234, doi:10.3389/fncel.2016.00234 (2016).

64 Dana, H. et al. Sensitive red protein calcium indicators for imaging neural activity. Elife 5, doi:10.7554/eLife.12727 (2016).

65 Namboodiri, V. M., Rodriguez-Romaguera, J. & Stuber, G. D. The habenula. Curr Biol 26, R873–R877, doi:10.1016/j.cub.2016.08.051 (2016).

66 Amo, R. et al. The habenulo-raphe serotonergic circuit encodes an aversive expectation value essential for adaptive active avoidance of danger. Neuron 84, 1034–1048, doi:10.1016/j.neuron.2014.10.035 (2014).

67 Turner, K. J. et al. Afferent Connectivity of the Zebrafish Habenulae. Front Neural Circuits 10, 30, doi:10.3389/fncir.2016.00030 (2016).

68 Bianco, I. H., Carl, M., Russell, C., Clarke, J. D. & Wilson, S. W. Brain asymmetry is encoded at the level of axon terminal morphology. Neural Dev 3, 9, doi:10.1186/1749-8104-3-9 (2008).

69 Agetsuma, M. et al. The habenula is crucial for experience-dependent modification of fear responses in zebrafish. Nat Neurosci 13, 1354–1356, doi:10.1038/nn.2654 (2010).

70 Amo, R. et al. Identification of the zebrafish ventral habenula as a homolog of the mammalian lateral habenula. J Neurosci 30, 1566–1574, doi:10.1523/JNEUROSCI.3690-09.2010 (2010).

71 Wang, R. Y. & Aghajanian, G. K. Physiological evidence for habenula as major link between forebrain and midbrain raphe. Science 197, 89–91, doi:10.1126/science.194312 (1977).

72 Chou, M. Y. et al. Social conflict resolution regulated by two dorsal habenular subregions in zebrafish. Science 352, 87–90, doi:10.1126/science.aac9508 (2016).

73 Turrini, L. et al. Fast multi-photon brain-wide volumetric imaging and photostimulation. SPIE BiOS 12384, doi:10.1117/12.2649824 (2023).

74 Hubert, A. et al. Random-access two-photon holographic optogenetic stimulation combined with brain-wide functional light-sheet imaging in larval zebrafish. European Conference on Biomedical Optics 12630, doi:10.1117/12.2671030 (2023).

75 Faini, G. et al. Ultrafast light targeting for high-throughput precise control of neuronal networks. Nat Commun 14, 1888, doi:10.1038/s41467-023-37416-w (2023).

76 Betzel, R. F. Organizing principles of whole-brain functional connectivity in zebrafish larvae. Netw Neurosci 4, 234–256, doi:10.1162/netn_a_00121 (2020).

77 Chicchi, L. et al. Reconstruction scheme for excitatory and inhibitory dynamics with quenched disorder: application to zebrafish imaging. J Comput Neurosci 49, 159–174, doi:10.1007/s10827-020-00774-1 (2021).

78 de Vito, G. et al. Two-photon high-speed light-sheet volumetric imaging of brain activity during sleep in zebrafish larvae. Neural Imaging and Sensing 2020 11226, doi:10.1117/12.2542285 (2020).

79 de Vito, G. et al. Two-photon light-sheet microscopy for high-speed whole-brain functional imaging of zebrafish physiology and pathology. Neurophotonics 11360, doi:10.1117/12.2560341 (2020).

80 Reddy, G. D. & Saggau, P. Fast three-dimensional laser scanning scheme using acousto-optic deflectors. J Biomed Opt 10, 064038, doi:10.1117/1.2141504 (2005).

81 Schneider, C. A., Rasband, W. S. & Eliceiri, K. W. NIH Image to ImageJ: 25 years of image analysis. Nat Methods 9, 671–675, doi:10.1038/nmeth.2089 (2012).

82 Bi, K. et al. Position of the prism in a dispersion-compensated acousto-optic deflector for multiphoton imaging. Appl Opt 45, 8560–8565, doi:10.1364/ao.45.008560 (2006).

83 Mullenbroich, M. C. et al. Bessel Beam Illumination Reduces Random and Systematic Errors in Quantitative Functional Studies Using Light-Sheet Microscopy. Front Cell Neurosci 12, 315, doi:10.3389/fncel.2018.00315 (2018).

84 ZFIN Feature: b4, <https://zfin.org/ZDB-ALT-980203-365.> (

85 Westerfield, M. The zebrafish book. A guide for the laboratory use of zebrafish (Danio rerio). 4th edn, (University of Oregon Press, 2000).

86 Turrini, L. et al. Multimodal Characterization of Seizures in Zebrafish Larvae. Biomedicines 10, 951, doi:10.3390/biomedicines10050951 (2022).

87 Johnsen, L. G., Skov, T., Houlberg, U. & Bro, R. An automated method for baseline correction, peak finding and peak grouping in chromatographic data. Analyst 138, 3502–3511, doi:10.1039/c3an36276k (2013).

88 Hellinger, E. Neue Begründung der Theorie quadratischer Formen von unendlichvielen Veränderlichen. Journal für die reine und angewandte Mathematik, doi:10.1515/crll.1909.136.210 (1909).

89 Ostrovsky, A., Cachero, S. & Jefferis, G. Clonal analysis of olfaction in Drosophila: image registration. Cold Spring Harb Protoc 2013, 347–349, doi:10.1101/pdb.prot071738 (2013).

90 Randlett, O. et al. Whole-brain activity mapping onto a zebrafish brain atlas. Nat Methods 12, 1039–1046, doi:10.1038/nmeth.3581 (2015).

91 Kunst, M. et al. A Cellular-Resolution Atlas of the Larval Zebrafish Brain. Neuron 103, 21–38 e25, doi:10.1016/j.neuron.2019.04.034 (2019).

92 Sorelli, M., Hutson, T. N., Iasemidis, L. & Bocchi, L. Linear and Nonlinear Directed Connectivity Analysis of the Cardio-Respiratory System in Type 1 Diabetes. Front Netw Physiol 2, 840829, doi:10.3389/fnetp.2022.840829 (2022).

93 Iatsenko, D., McClintock, P. V. E. & Stefanovska, A. Linear and synchrosqueezed time-frequency representations revisited: Overview, standards of use, resolution, reconstruction, concentration, and algorithms. Digit Signal Process 42, 1–26, doi:10.1016/j.dsp.2015.03.004 (2015).

